# Microtubule occupancy at kinetochores links checkpoint silencing with mitotic memory

**DOI:** 10.64898/2026.01.26.701783

**Authors:** Joana Soares-de-Oliveira, Naoyuki Okada, Tobias Kletter, Elias S. Weiler, António J. Pereira, Helder Maiato

## Abstract

The spindle assembly checkpoint (SAC) promotes faithful chromosome segregation by delaying mitosis until all kinetochores attach to spindle microtubules. Paradoxically, a p53-dependent memory mechanism—the mitotic stopwatch” —blocks daughter cell proliferation after unusually prolonged mitoses. How the SAC coordinates with the mitotic stopwatch remains unknown. Here, we found that microtubule occupancy at kinetochores is a cornerstone linking SAC silencing with mitotic memory. By combining live-cell with super-resolution microscopy, FRAP, laser microsurgery, biochemistry and molecular perturbations in Indian muntjac fibroblasts, we show that SAC silencing at kinetochores is gradual, non-uniform and confined to highly-localized microtubule attachments. Augmin promotes timely SAC silencing with high microtubule occupancy at kinetochores, whereas MPS1/CDK1 inhibition bypasses this requirement. Conversely, low microtubule occupancy delays SAC silencing, increases segregation errors and blocks daughter cell proliferation due to mitotic stopwatch surveillance. Thus, timely SAC silencing with high microtubule occupancy avoids “bad memories” of mitosis to allow daughter cell proliferation.

## Introduction

Faithful chromosome segregation during mitosis is essential for tissue homeostasis. Errors in chromosome segregation may lead to aneuploidy compromising cell viability, or drive chromosomal instability, a hallmark of human cancers implicated in tumor evolution, metastasis, therapy resistance, and reduced patient survival ^1^. To promote faithful chromosome segregation during mitosis, the spindle assembly checkpoint (SAC) delays anaphase until all kinetochores attach to mitotic spindle microtubules ^2^. At the root of the SAC signaling cascade, a MAD1/MAD2 complex recruited to unattached kinetochores initiates a catalytic process that activates free cytosolic MAD2 to bind CDC20, BUBR1 and BUB3, producing a diffusible Mitotic Checkpoint Complex (MCC). The MCC inhibits the Anaphase Promoting Complex/Cyclosome (APC/C), an E3-ubiquitin ligase that otherwise targets Cyclin B1 and Securin for degradation by the proteasome (reviewed in ^3–5^). This releases the cysteine-protease Separase to cleave the Cohesin molecules that hold sister-chromatids together, thereby triggering their initial separation in anaphase, while preventing SAC reactivation during anaphase ^6,7^.

Upon kinetochore attachment to microtubules, SAC strength gradually decreases as result of the removal of MAD1/MAD2, eventually halting the catalytic cascade underlying MCC production ^8,9^. The removal of MAD1/MAD2 from kinetochores during SAC silencing is mediated by PP1 and PP2A phosphatases, which are recruited by the outer kinetochore component KNL1 to oppose the action of key mitotic kinases, such as CDK1 and MPS1^10–15^. As microtubules establish end-on attachments, MPS1 is outcompeted from NDC80 complexes at kinetochores^16,17^, and the MAD1/MAD2 complex is removed by dynein-mediated stripping along with the disassembly of the fibrous corona ^18–22^. Pioneering correlative light and electron microscopy work in rat kangaroo PtK1 cells showed that their kinetochores bind more microtubules in anaphase than in metaphase, indicating that SAC silencing during normal mitosis can occur at ∼85% microtubule occupancy at kinetochores ^23^. Classic micromanipulation experiments in grasshopper spermatocytes further indicated that weakly attached kinetochores with less than 40% microtubule occupancy show greatly diminished MAD1/MAD2 recruitment when compared with fully unattached kinetochores, delaying but not preventing anaphase onset ^24^. Indeed, in this system, high microtubule occupancy was found to be necessary for the reliable and complete removal of MAD1/MAD2 from kinetochores ^24^. In contrast, two recent studies in human HeLa cells overexpressing individual or combinations of Hec1 phosphomimetic mutants to create intermediate average microtubule attachment states at kinetochores estimated that 20–50% of the normal metaphase microtubule occupancy is sufficient to trigger complete and uniform removal of MAD1/MAD2 from kinetochores ^25,26^. However, this switch-like model remains difficult to reconcile with the finding that the rate of MAD1/MAD2 loss from kinetochores increases with microtubule occupancy, with SAC silencing under conditions of low microtubule occupancy resulting in extensive mitotic delays ^24–26^. A third study in human RPE1 cells used a microtubule depolymerizing drug to reduce microtubule occupancy at kinetochores ^27^. Based on direct microtubule counting by electron microscopy, this study showed that ∼70% microtubule occupancy at metaphase kinetochores is sufficient to silence the SAC, with less than 5 min delay relative to controls, but at the price of increasing the incidence of lagging chromosomes during anaphase ^27^. Thus, although cells appear to be able to silence the SAC with low microtubule occupancy, there may be selective pressure for high microtubule occupancy at kinetochores before initiating anaphase, ensuring proper chromosome segregation and avoiding significant mitotic delays ^27,28^. This hypothesis gains weight in light of the recently uncovered “mitotic stopwatch” complex formed by 53BP1 and the deubiquitinase USP28 that stabilizes p53 in response to even moderate delays in mitosis, leading to subsequent cell cycle arrest or death in G1, independently of DNA damage ^29–36^. However, how SAC silencing is coordinated with the mitotic stopwatch to control daughter cell proliferation remains unknown.

Here we set to investigate how the SAC responds to highly-localized microtubule attachments at kinetochores, while testing whether the extent of microtubule occupancy underlying SAC silencing impacts daughter cell proliferation. Overall, our data support a gradual, non-uniform (i.e. confined to highly-localized microtubule attachments), SAC-silencing mechanism that is assisted by Augmin to promote high microtubule occupancy at kinetochores for timely and faithful chromosome segregation. Importantly, if cells ultimately silence the SAC after experiencing significant mitotic delays due to low microtubule occupancy at kinetochores, the resulting daughter cells are bookmarked by the mitotic stopwatch to prevent their proliferation and the potential propagation of segregation errors.

## Results

### MAD1 is gradually removed from Indian muntjac kinetochores in a microtubule-dependent manner

According to the switch-like SAC-silencing model, kinetochores respond as a single unit to silence the SAC after just a few microtubule attachments are established ^25,26^. However, because human kinetochores cannot be resolved by conventional light microscopy, this postulate has so far been impossible to test experimentally. To overcome this limitation, we took advantage of the unique cytological features of female Indian muntjac fibroblasts, which carry only six chromosomes with naturally “super-resolved” kinetochores that arose by tandem and centric fusions of smaller ancestral chromosomes ^37–43^. As a first step towards investigating SAC silencing in this system, we determined the time between MAD1 disappearance from kinetochores and anaphase onset by tracking individual kinetochores in live, hTERT-immortalized, female Indian muntjac fibroblasts stably expressing mScarlet-CENP-A (a constitutive kinetochore marker) and Venus-MAD1 ^8^. We found that MAD1 signal at kinetochores became undetectable ∼20 min before anaphase onset (Fig. S1A, B), in agreement with previous reports in other vertebrate cell lines ^44,45^ and consistent with the temporal framework of SAC silencing upon laser-mediated ablation of the last unattached kinetochore ^2^.

To determine the kinetics of SAC silencing in response to microtubule attachments at individual kinetochores in live Indian muntjac fibroblasts, we combined stable expression of fluorescent Venus-MAD1 with carefully titrated SiR-tubulin ^41^ under slight cell compression to improve visualization ^42^. As a control, cytoplasmic Venus-MAD1 signal was simultaneously measured, and photobleaching was found to be negligible throughout the course of the experiment (Fig. S1C). MAD1 decayed gradually at individual kinetochores as chromosomes bi-oriented and aligned at the equator during prometaphase (Figure 1A, C and Video S1), with a rate of decay that was well fit to a single exponential (Figure 1E and Fig. S2A), in agreement with previous reports in PtK2 cells ^46^. Noteworthy, pole-proximal chromosomes aligned last and did so tangentially along the curvature of the spindle edge. Moreover, their leading kinetochores were enriched with MAD1 from the onset of congression, with the signal gradually decreasing as chromosomes approached the equator (Figure 1A, A’, A’’ and Video S1). Importantly, MAD1 typically decayed first on trailing kinetochores, likely reflecting differences in the timing or efficiency in the formation of stable end-on microtubule attachments compared with leading kinetochores. (Figure 1A, A’, A’’ and Video S1). These results are fully consistent with congression models where pole-proximal, mono-oriented chromosomes use plus-end-directed motors on the leading kinetochore to initiate lateral gliding along pre-existing spindle microtubules towards the equator ^46–49^, while at odds with models where stabilization of end-on attachments and bi-orientation take place before pole-proximal chromosomes initiate congression ^50^.

**Figure 1.**
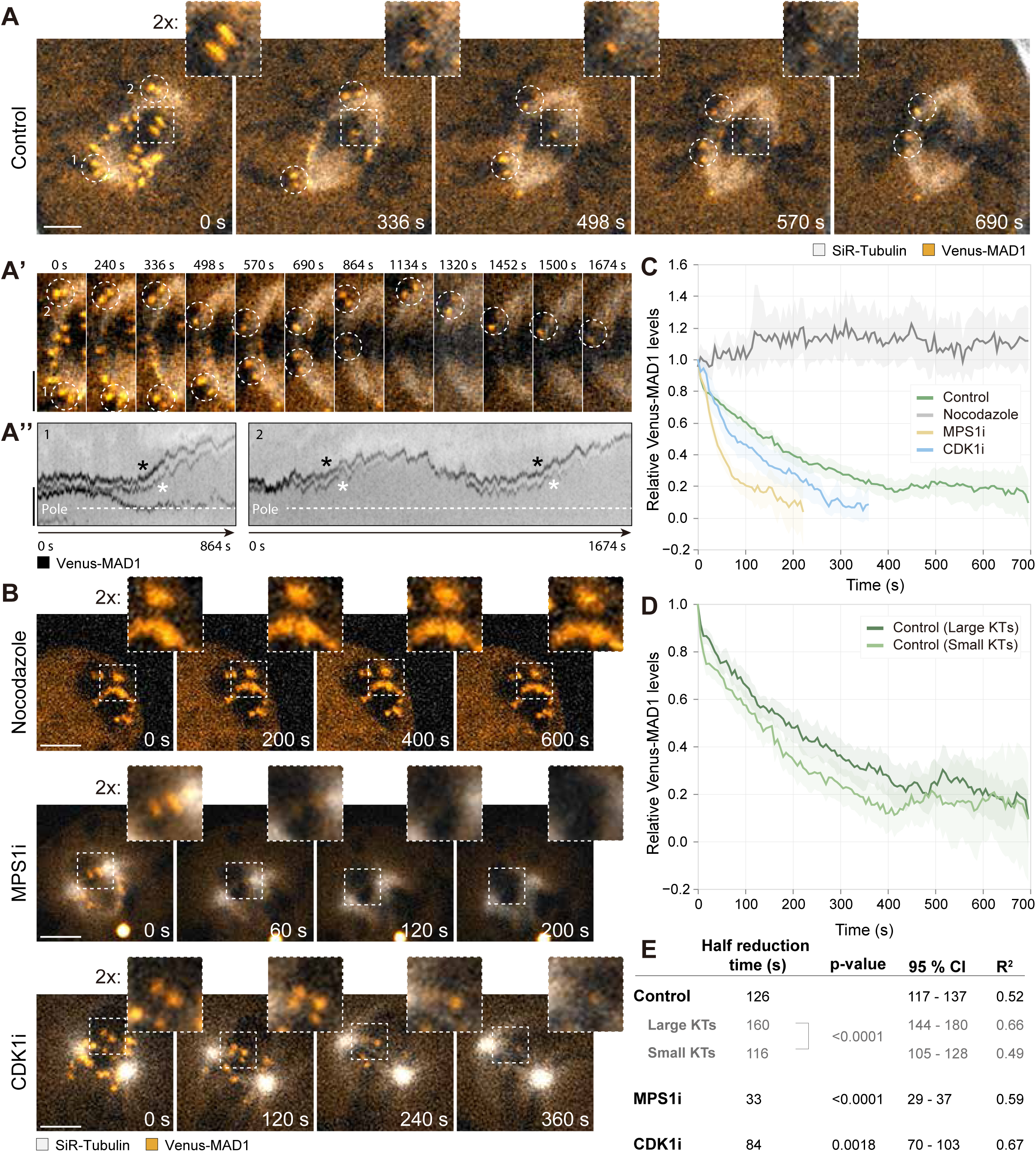
MAD1 is gradually removed from the kinetochore in a microtubule-and mitotic kinases-dependent manner. (A) Representative live-cell recording images of Indian muntjac fibroblasts expressing Venus-MAD1 (orange) and with SiR-tubulin labeled microtubules (grey). Scale bar: 5 µm. Insets highlight an early congressing kinetochore pair (dashed square) and dashed circles show polar kinetochores/chromosomes. Scale bar: 5 µm. (A’) Insets focusing on 1 half-spindle to highlight the congression of polar chromosomes (dashed circles) from A. Scale bar: 5 µm. (A’’) Kymographs depicting Venus-MAD1 signal of the 2 polar chromosomes shown in A’. The closest pole position is shown by the dashed line. Leading (black asterisk) and trailing kinetochore (white asterisk) and indicated for each pair. Scale bar: 5 µm. (B) Representative live-cell recording images of Indian muntjac fibroblasts acutely treated with Nocodazole, MPS1 and CDK1 inhibitors. Venus-MAD1 (orange) and SiR-tubulin labeled microtubules (grey). Insets highlight a kinetochore pair. Scale bar: 5 µm. (C) Normalized Venus-MAD1 fluorescence decay IM cells untreated (control) or treated with Nocodazole, MPS1 and CDK1 inhibitors. Data represented as mean ± 95% confidence interval (CI; Control n = 53 kinetochores, 7 cells, green; Nocodazole n = 31 kinetochores, 8 cells, grey; MPS1i n = 63 kinetochores, 12 cells, yellow; CDK1i n = 18 kinetochores, 10 cells, blue; data pooled from at least five independent experiments per condition). (D) Normalized Venus-MAD1 fluorescence decay in large or small kinetochores from untreated cells. Data are represented as mean ± 95% CI. (Large KTs n = 19 kinetochores, 7 cells, 5 independent experiments, dark green; small KTs n = 34 kinetochores, 5 cells, 4 independent experiments, light green). (E) Summary table of the parameters extracted from a single exponential fitting of Venus-MAD1 fluorescence decay across the condition shown in C and D. P-values were calculated using extra sum-of-squares *F*-test comparing the rate of MAD1 decay (K, derived from global fit) in untreated vs treated cells or large vs small kinetochores.

To test whether gradual MAD1 decay at kinetochores during mitosis depends on microtubule occupancy, we monitored MAD1 levels in the presence of 1 µM nocodazole, which depolymerized all spindle microtubules. As expected, under these conditions, MAD1 localized on every kinetochore for over 20 min, without any measurable decay (Figure 1B, C, Fig. S2A and Video S2). These data indicate that gradual SAC silencing during mitosis depends on the establishment of microtubule attachments at kinetochores.

### MAD1 is rapidly removed from Indian muntjac kinetochores upon MPS1 or CDK1 inactivation, regardless of microtubule occupancy

MPS1 and CDK1 kinases are critical to maintain SAC signaling by counteracting the activities of PP1 and PP2A phosphatases required for mitotic exit ^10–15^. Therefore, we next investigated the kinetics of MAD1 decay at kinetochores after acute MPS1 or CDK1 inactivation with MPS1-IN-1 ^51^ or RO 3306 ^52^, respectively. We found that MAD1 decayed more abruptly from kinetochores after MPS1 or CDK1 inhibition, regardless of microtubule occupancy, reaching 50% of the initial levels 3.8x or 1.5x faster than during normal mitosis, respectively (Figure 1B, C, E, Fig. S2A and Videos S3, S4). These data indicate that inactivation of MPS1 or CDK1 can bypass the requirement of a given microtubule occupancy at kinetochores to silence the SAC.

### MAD1 removal depends on kinetochore size

A prediction from switch-like models that favor independent microtubule binding at kinetochores and where a given percentage of occupancy is key to uniformly silence the SAC ^25,26^ is that small or large kinetochores would simultaneously reach the same percentage of occupancy (and, consequently, have similar MAD1 half reduction times), despite binding a different total number of microtubules. However, despite ample evidence of kinetochore size differences in humans ^53–61^, this prediction is currently impossible to test experimentally in human cells for the same reasons evoked before. In contrast, the large kinetochore from chromosome 3+X in female Indian muntjac cells can be unequivocally distinguished from those on the smaller chromosomes 1 and 2, allowing us to track MAD1 decay as a function of kinetochore size ^38,40,41^. Surprisingly, we found that MAD1 signal on the large kinetochores of chromosome 3+X requires ∼38% more time to become undetectable, when compared with smaller kinetochores on chromosomes 1 and 2 (Figure 1D, E, and Fig. S2A). These findings suggest that switch-like models do not fully account for how the SAC is normally silenced.

### MAD1 is immobile within kinetochores

In order to obtain a deeper understanding of the mechanisms underlying SAC silencing we investigated MAD1 dynamics within kinetochores. Two possible scenarios could be envisioned: 1) if MAD1 is mobile within kinetochores, few microtubule attachments might be sufficient to “extinguish” all MAD1, for example by Dynein-mediated stripping along microtubules ^18,19^ (Figure 2A); 2) if MAD1 is immobile within kinetochores, Dynein-mediated stripping along microtubules would be confined to the sites or domains where microtubules are attached (Figure 2A). To distinguish between these two scenarios, we photobleached Venus-MAD1 on a fraction (up to 50%) of the large kinetochore from chromosome 3+X in the absence of microtubules. Clear predictions about MAD1 mobility within kinetochores could be made, depending on the FRAP profile. If MAD1 is mobile within kinetochores, the photobleached fraction of the kinetochore is expected to recover due to migration of fluorescent Venus-MAD1 molecules from the unbleached fraction, resulting in the convergence of signals to an intermediate and uniform level along the entire kinetochore (Figure 2A’). In contrast, if MAD1 is immobile within kinetochores, the photobleached fraction of the kinetochore would be expected to only recover fluorescent Venus-MAD1 molecules that are exchanging with the cytoplasmic pool, while the unbleached fraction would remain stable over time (Figure 2A’). We were successful in partially photobleaching Venus-MAD1 in a fraction of the kinetochore, with minimal/unintended photobleaching of Venus-MAD1 on the remaining fraction (Figure 2B-B’’ and Video S5). Importantly, after photobleaching of the intended fraction of the kinetochore, Venus-MAD1 fluorescence recovered up to ∼50%, with a half recovery time of 2-6 seconds, while remaining stable (or increasing slightly; see ahead) over time in the unbleached fraction (Figure 2B-B’’, D, E and Video S5). As a control, we photobleached Venus-MAD1 on entire kinetochores, which resulted in identical recovery parameters when compared to partial photobleaching, and in line with previous measurements in other mammalian systems ^44,45^ (Figure 2C-C’’, D, E and Video S6). Because fluorescence recovery of the mobile fraction of Venus-MAD1 after photobleaching entire kinetochores is exclusively due to exchange with the cytoplasmic pool ^44,45^, we concluded that this pool also accounts for the observed fluorescence recovery of Venus-MAD1 on partially photobleached kinetochores. Taken together, these data indicate that MAD1 is essentially immobile within kinetochores, suggesting that it can only be removed from spatially confined regions upon the establishment of highly-localized microtubule attachments or by a microtubule-independent SAC-silencing mechanism, such as the one involving CDK1 or MPS1 inactivation.

**Figure 2.**
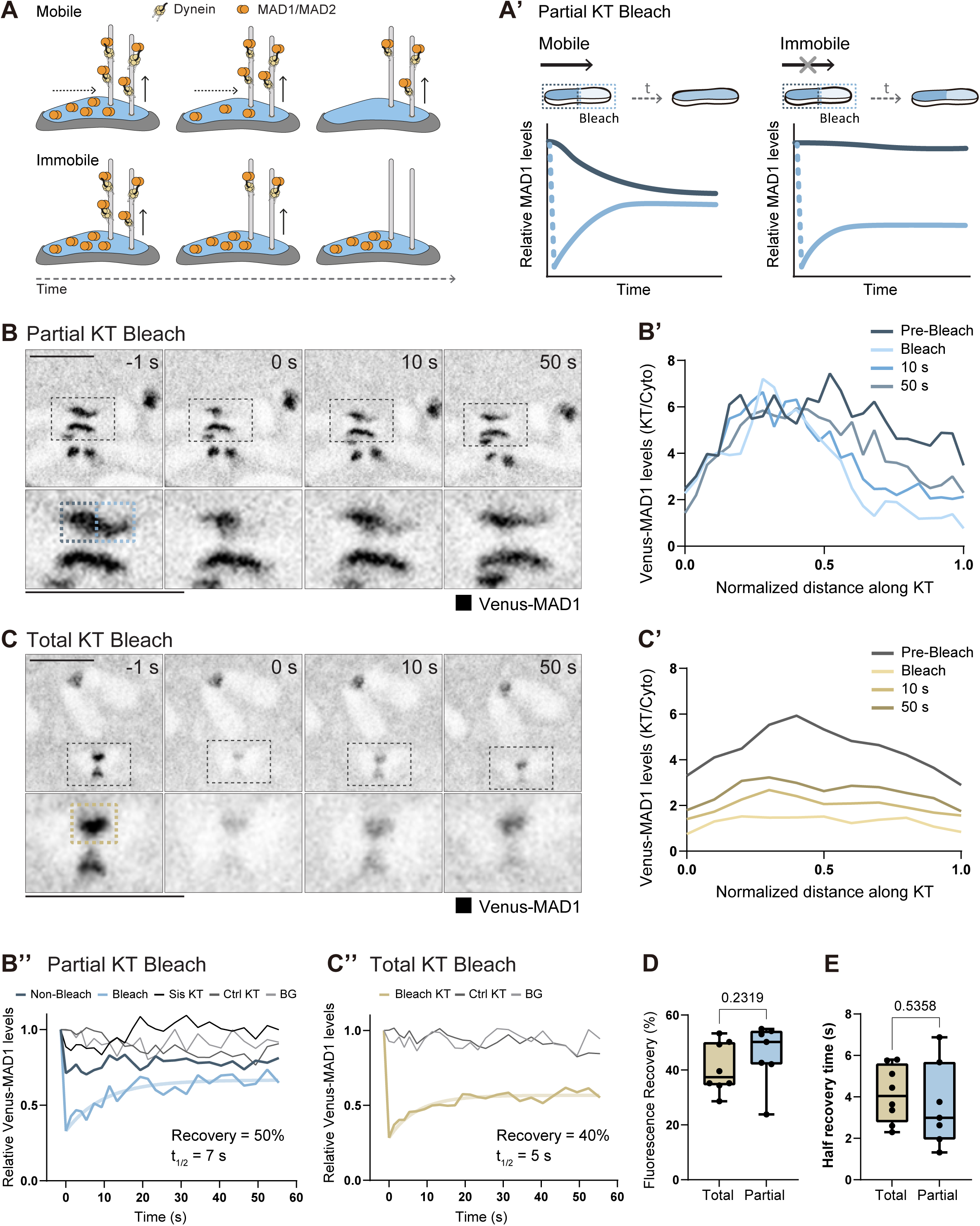
MAD1 is immobile within kinetochores (A) Schematic representation of Dynein-mediated MAD1/MAD2 complex removal from kinetochores. (A’) Predicted curves of MAD1 fluorescence recovery after partial kinetochore bleaching assuming MAD1 to be mobile (left) or immobile (right) within the kinetochore. Lines represent the fluorescence levels from the region of the kinetochore that was bleached (light blue) or left unperturbed (dark blue). (B), (C) Representative images of Indian muntjac cells pre-and post-bleach of part (B) or an entire kinetochore (C). Insets: kinetochore pair that was perturbed. Dashed line boxes indicate the ablated kinetochore region (partial bleach KT: non-bleached region - dark blue; bleached region – blue; total bleach KT - yellow). (B’), (C’) Venus-Mad1 signal intensity profile along the kinetochore at pre-bleach and at selected times post-bleach for kinetochores shown in B and C, respectively. (B’’), (C’’) Normalized Venus-Mad1 fluorescence recovery curves for cells shown in B and C, respectively (Dark blue line non-bleached region; light blue bleached region; yellow line totally bleached kinetochore; black and dark grey lines unperturbed kinetochores from the bleach or a control kinetochore pair, respectively; light grey cytoplasm; from fit curves are shown by the semi-translucid line). (D) Kinetochore Venus-MAD1 fluorescence recovery after partial or total kinetochore bleach (Partial KT bleach n = 7; total KT bleach n = 8 from at least two independent experiments). Dots show individual kinetochores. P = 0.2319, Mann-Whitney test (two-tailed). (E) Half recovery times of kinetochore Venus-MAD1 fluorescence after partial or total kinetochore bleach. Dots show individual kinetochores. P = 0.5358, Mann-Whitney test (two-tailed).

### MAD1 removal along individual kinetochores is non-uniform and confined to highly-localized microtubule attachments

The conclusion from our FRAP experiments implies that, in a normal mitosis, there may be low microtubule occupancy states in which MAD1 removal is limited to regions where microtubule attachments are established. To test this prediction, we monitored Venus-MAD1 decay along the entire kinetochore length as microtubules attach during mitosis. Near-simultaneous tracking of individual kinetochores, MAD1 and microtubules in living cells revealed a non-uniform decay of MAD1 along kinetochores (Fig. S2B-B’). To further dissect the relationship between non-uniform MAD1 decay and microtubule occupancy along kinetochores, we used super-resolution CH-STED microscopy ^62^ and 3D image rendering to inspect (fixed) late prometaphase cells with only partial MAD1 signal along kinetochores (Figure 3A, B and Fig. S3A, B). We found clear examples where fractions of the kinetochore had end-on attached microtubules with no detectable MAD1, while other fractions were devoid of microtubules and clearly enriched for MAD1 (Figure 3A, B and Fig. S3A, B). A line scanning profile along the entire length of several such individual kinetochores confirmed a negative correlation between MAD1 signal and microtubule density in 15/21 (71.4%) of the cases (Figure 3C, D and Fig. S3A’, B’).

**Figure 3.**
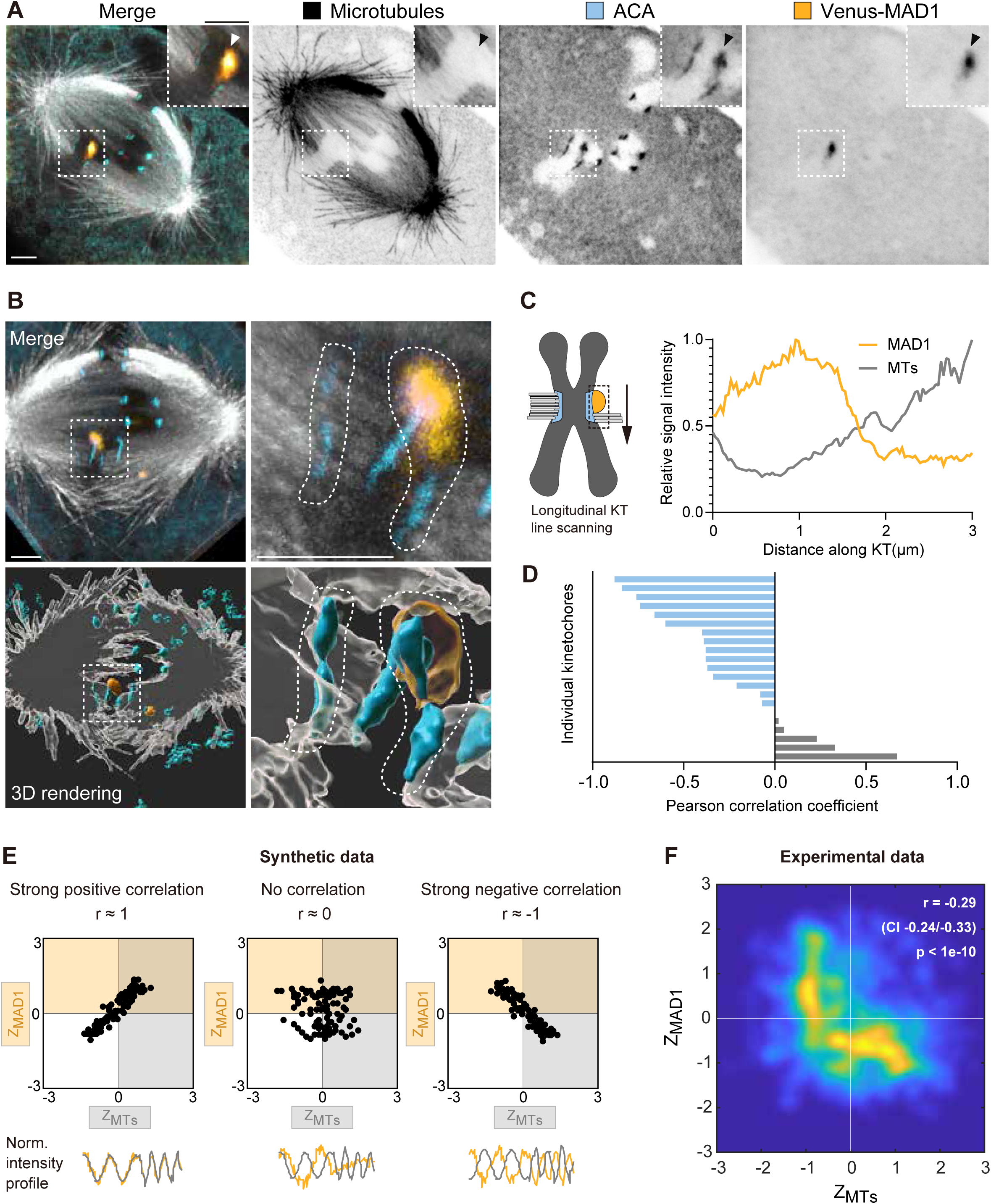
MAD1 removal along individual kinetochores correlates with local microtubule density (A) Representative CH-STED image of an Indian muntjac fibroblasts with partially attached kinetochores. Microtubules (grey; STED), centromeres (ACA; blue; STED) and Venus-MAD1 (orange; confocal). Insets: partially attached kinetochore. Scale bar: 2 µm. (B) Three-dimensional reconstruction of the cell shown in A (different perspective). Inset: partially attached kinetochore. The kinetochore pair of interest is traced by a dashed line. Microtubules (grey), centromeres (ACA; blue) and Venus-MAD1 (orange). Scale bars, Left: 3 µm; Right: 1 µm. (C) Max normalized Venus-Mad1 (orange) and tubulin (microtubules; grey) signal intensity profile along the longitudinal axis of the kinetochore highlighted in A (arrowhead). (D) Pearson correlation coefficient between MAD1 and microtubules signal intensity along the kinetochore. Bars show individual partially attached kinetochores (n=21 kinetochores, 7 independent experiments). E) Illustrative/simulated distribution of kinetochore sub-regions showing positive, negative and no correlation between MAD1 (y-axis) and microtubules (x-axis) intensity z-scores. Dots represent distinct positions along the “kinetochore” shown below the graphs. NE and SW quadrants contribute positively, while NW and SE quadrants contribute negatively to the Pearson correlation coefficient (r). (F) Correlation map of MAD1 and microtubule intensity z-scores for the kinetochores shown in D. The underlying scatterplot contains data from multiple kinetochores, with each dot representing a sub-kinetochore region. For visualization, the scatterplot was converted into a regional density map. P-value was calculated with Student t-test for zero correlation (two-sided).

To improve the resolution of our analyses we divided the longitudinal axis of partially attached kinetochores into multiple subsections and quantified MAD1 and microtubule intensity at each subsection (Figure 3E). We found a weak, yet highly significant, negative correlation between MAD1 and microtubule signal intensity along kinetochores (Figure 3F and Fig. S3C). Of note, the resulting L-shape curve indicates that while high microtubule signal is a good predictor of the absence of MAD1 at kinetochores, the opposite is not the case, suggesting that there are regions at kinetochores without microtubules that have no detectable MAD1, consistent with models evoking cooperative microtubule binding at kinetochores ^63,64^. Overall, these data show that SAC silencing at kinetochores is non-uniform and confined to highly-localized microtubule attachments.

### Localized kinetochore perturbations result in a local SAC response to microtubule attachments/detachments

To directly test how the SAC responds to localized microtubule attachments/detachments, we used laser microsurgery to partially ablate individual metaphase kinetochores (and, consequently, microtubules that were attached to the ablated region) upon SAC silencing in live Indian muntjac fibroblasts (Figure 4A). These cells stably expressed GFP-CENTRIN-1 to mark spindle pole positions, 2xGFP-CENP-A to define the kinetochore target, and mScarlet-MAD1 to monitor SAC status before and after surgery (Figure 4B and Video S7). We found that, within few minutes after partial kinetochore ablation, the unperturbed sister kinetochore typically bent towards the attached spindle pole without any detectable MAD1 (Figure 4A-C and Video S7). However, ∼5 min after bending, MAD1 was often recruited to the tip of the bent unperturbed sister kinetochore, presumably due to highly-localized microtubule detachments (Figure 4A-D and Video S7). In other cases, MAD1 was recruited to a more central region of the bent unperturbed sister kinetochore (Fig. S4A, A’, B). Strikingly, in all cases MAD1 accumulation was highly localized and distinct from the uniform accumulation of MAD1 on fully unattached kinetochores resulting from nocodazole treatment (Figure 4B, D, E). Therefore, local kinetochore perturbations result in a highly-localized SAC response.

**Figure 4.**
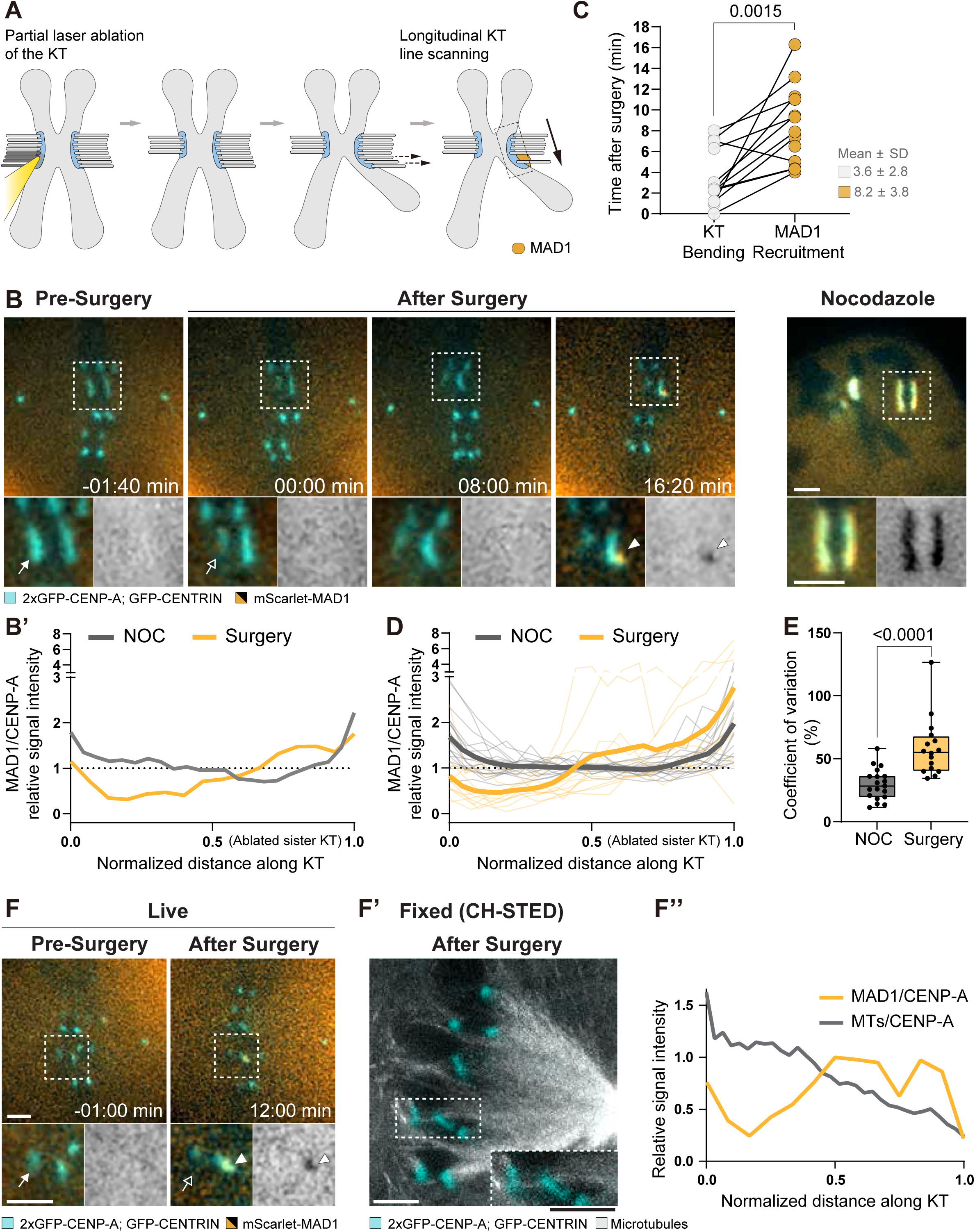
Local kinetochore perturbation results in a localized SAC response that inversely correlates with microtubule occupancy (A) Schematic summary of the kinetochore response to partial laser microsurgery ablation. (B) Representative live-recording images of Indian muntjac cells expressing 2x-GFP-CENP-A and GFP-CENTRIN-1 (both in blue) and mScarlet-MAD1 (orange/ grey scale) upon partial kinetochore surgery or acute Nocodazole treatment. Insets: kinetochore pair that was perturbed. Arrows point to the ablated kinetochore region, pre-and post-surgery (solid and hollow arrow, respectively). Arrowheads show MAD1 recruitment. Scale bar: 2 µm. (B’) Max normalized line intensity profile plot of MAD1/CENP-A signal ratio along the longitudinal axis of the kinetochores highlighted in A (surgery cell in orange; nocodazole cell in grey). (C) Dot plot showing the time from surgery to observable deformation (bending; n= 12 kinetochores) and MAD1 signal re-appearance (n = 14 kinetochores) at the sister kinetochore in cells where SAC recruitment was observed. Each dot represents an individual kinetochore from a different cell, recorded during 13 independent experiments. P <0.0015, Wilcoxon matched-pairs signed rank test. (D) Max normalized MAD1/CENP-A ratio intensity profile along the longitudinal axis of kinetochores after partial surgery of their sisters (orange) or upon acute Nocodazole treatment (NOC; grey). Thin lines show individual kinetochores and thick lines show average curves (surgery n = 11 kinetochores/cells, 10 independent experiments; Nocodazole n = 19 kinetochores/cells, 2 independent experiments). The graph depicts a cohort of cells that recruit MAD1 to the proximal tip of the kinetochore, relative to the surgery region (n = 11/16 MAD1 recruiting kinetochores). (E) Coefficient of variation of MAD1/CENP-A signal ratio along kinetochores after partial surgery of their sisters. Dots represent individual kinetochores pooled from all cells where MAD1 recruitment was observed (surgery n = 16 cells, 14 independent experiments; Nocodazole n = 19 cells, 2 independent experiments). P <0.0001, Mann-Whitney test. (F), (F’) Representative images from correlative live cell spinning disk (F) and fixed CH-STED microscopy (F’). Indian muntjac cells expressing 2x-GFP-CENP-A and GFP-CENTRIN-1 (both in blue) and mScarlet-MAD1 (orange/grey scale) were fixed after partial kinetochore surgery and stained to visualize microtubules (grey). Insets: kinetochore pair that was perturbed. Arrows point to the ablated kinetochore region, pre-and post-surgery (solid and hollow arrow, respectively). Arrowheads show MAD1 recruitment. Scale bar: 2 µm. (F’’) Max normalized line intensity profile plot of MAD1/CENP-A (live cell data just before fixation, orange) and microtubules/CENP-A (fixed cell data, grey) signal ratio along the longitudinal axis of the kinetochore highlighted in F and F’.

To assist us in the interpretation of the laser microsurgery data and determine whether the SAC was responding to microtubule detachments in the bent unperturbed sister kinetochore, we performed correlative super-resolution CH-STED microscopy after fixation of cells that showed MAD1 recruitment upon partial sister kinetochore laser ablation by live imaging (Figure 4F, F’ and Fig. S4C, C’). We found that MAD1 accumulation, either at the tip or at a more central region of the unperturbed sister kinetochore, negatively correlated with microtubule density (Figure 4F’, F’’ and Fig. S4C’, C’’). We reasoned that MAD1 accumulation near the tip of the unperturbed kinetochore may result from highly localized microtubule detachments due to lack of tension in the partially ablated sister. In other cases, localized forces applied to both ends of the unperturbed kinetochore may generate enough tension along its length to allow a few microtubules to remain stably attached near the tip, resulting in more central accumulation of MAD1. These results demonstrate that the SAC responds locally to highly localized microtubule attachment/detachment events along kinetochores.

### Low microtubule occupancy at kinetochores after Augmin depletion delays MAD1 removal and SAC silencing

Next, we sought to investigate the cellular consequences of low microtubule occupancy at kinetochores. To do so we monitored SAC silencing after RNAi-mediated depletion of HAUS6, a subunit of the Augmin complex required for K-fiber maturation ^43,65^. We found that MAD1 decay from kinetochores was slowed-down in HAUS6-depleted cells, as revealed by ∼50% increase in the half reduction time of MAD1 signal at kinetochores relative to mock-depleted controls (Figure 5A-C, Figure S2A and Videos S8, S9). Moreover, even when HAUS6-depleted cells were treated with the proteasome inhibitor MG132 for 1h to provide more time for K-fiber maturation, many more (66% vs. 12%) cells had at least one MAD1-positive kinetochore and overall higher MAD1 levels at kinetochores when compared with control metaphase cells (but lower than prometaphase levels) (Figure 5D, E). Closer inspection of mock-and HAUS6-depleted fixed cells with CH-STED super-resolution microscopy revealed that, while kinetochores in control mock-depleted metaphase cells treated with MG132 were abundantly occupied with microtubules and without detectable MAD1 signal, in HAUS6-depleted cells treated with MG132 MAD1 signal could still be detected along the kinetochores, despite interspersed end-on microtubule attachments (Figure 5F). However, unlike in control cells with clear partially attached kinetochores that showed a negative correlation between MAD1 and microtubule signal (r =-0.29, p<1e-10; Figure S5E), in HAUS6-depleted cells this correlation was lost (r =-0.06, p=0.14; Figure S5D, E and Figure S6C), suggesting that the resulting perturbations of microtubule occupancy were beyond our resolution capacity (Fig. S5A-C and Fig. S6A-B’). Overall, these data are in line with previous findings in human cells ^25,26^ indicating that low microtubule occupancy at kinetochores delays MAD1 removal from kinetochores and consequently SAC silencing.

**Figure 5.**
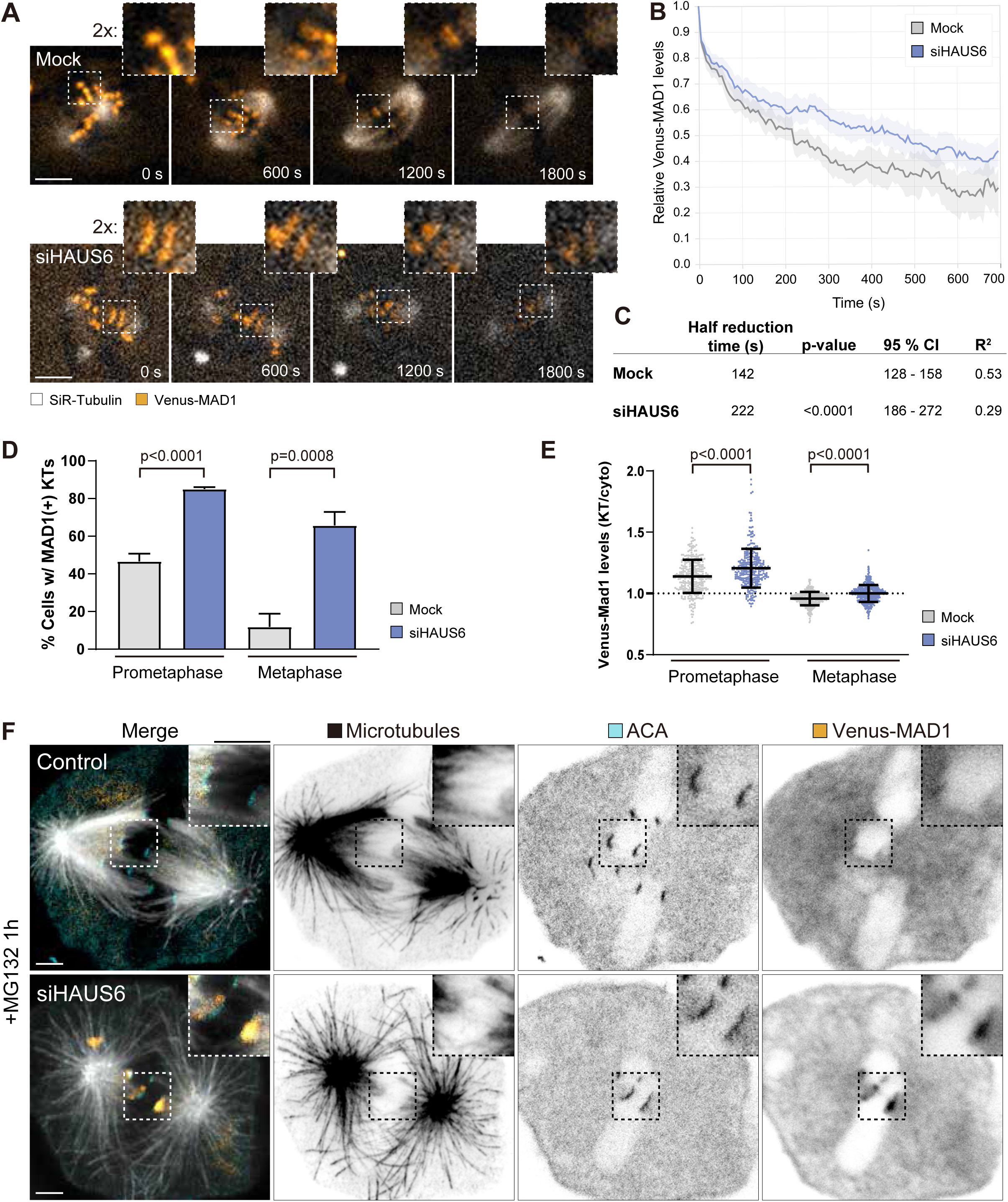
Partial microtubule occupancy at kinetochores after Augmin depletion delays MAD1 removal and SAC silencing (A) Representative live-cell recording images of Indian muntjac fibroblasts mock or HAUS6 siRNA treated. Venus-MAD1 (orange) and SiR-tubulin labeled microtubules (grey). Scale bar: 5 µm.Insets highlight a kinetochore pair. Asterisks indicate the two kinetochores from the pair. Scale bar: 5 µm. (B) Normalized Venus-MAD1 fluorescence decay from A. Data from at least five independent experiments are represented as mean ± 95% confidence interval (Mock n = 48 kinetochores, 10 cells, grey; siHAUS6 n = 47 kinetochores, 10 cells, blue). (C) Summary table of the parameters extracted from a single exponential fitting of Venus-MAD1 fluorescence decay across the conditions shown in B. P-values were calculated using extra sum-of-squares *F*-test comparing the rate of MAD1 decay (K, derived from global fit) in mock vs siHAUS6 treated cells. (D) Percentage of cells with at least one MAD1 positive kinetochore in mock (grey) or siHAUS6 (blue) treated cells after 1h of MG132 arrest. Bars show mean, error bars show s.d. from three independent experiments (n ≥ 293 cells per condition). p <0.0001, prometaphase; p = 0.0008, metaphase; unpaired t test (two-sided). (E) Dot plot showing the ratio between kinetochore and cytoplasmatic Venus-MAD1 signal intensity in mock (grey) or siHAUS6 (blue) treated cells fixed after 1 h of MG132 arrest. Dots show individual kinetochores, bars show mean, error bars show s.d. from three independent experiments (n≥309 kinetochores, ≥17 cells per condition). p <0.0001, Mann-Whitney test (two-sided). (F) Representative CH-STED image of an Indian muntjac control or HAUS6 depleted cells treated with MG132 for 1 h. Microtubules (grey; STED), centromeres (ACA; blue; STED) and Venus-MAD1 (orange; confocal). Inset: full (control) or partially (siHAUS6) attached kinetochore pair. Scale bar: 2 µm.

### Low microtubule occupancy delays SAC silencing, increases segregation errors, and blocks daughter cell proliferation through a conserved stopwatch mechanism

To further characterize the cellular consequences of low microtubule occupancy at kinetochores, we tracked HAUS6-depleted cells over several days by live-cell microscopy and determined their fates (Figure 6A). Mitoses in HAUS6-depleted cells were significantly delayed and error-prone, proportional to the extent of HAUS6 depletion over time (Figure 6B, 6C-6C’’). When inspecting the outcomes resulting from mitotic delays in HAUS6-depleted cells more closely, we found three regimens. The majority of cells with mitotic durations below 200 minutes divided without segregation errors or spindle defects (Figure 6C). However, after a delay of approx. 200 minutes, virtually all cells divided with mitotic defects (Figure 6C), pointing towards persistently immature K-fibers at anaphase onset^27^. Strikingly, most cells that were stuck in mitosis for longer than 600 minutes ultimately died in mitosis (Figure 6C).

**Figure 6.**
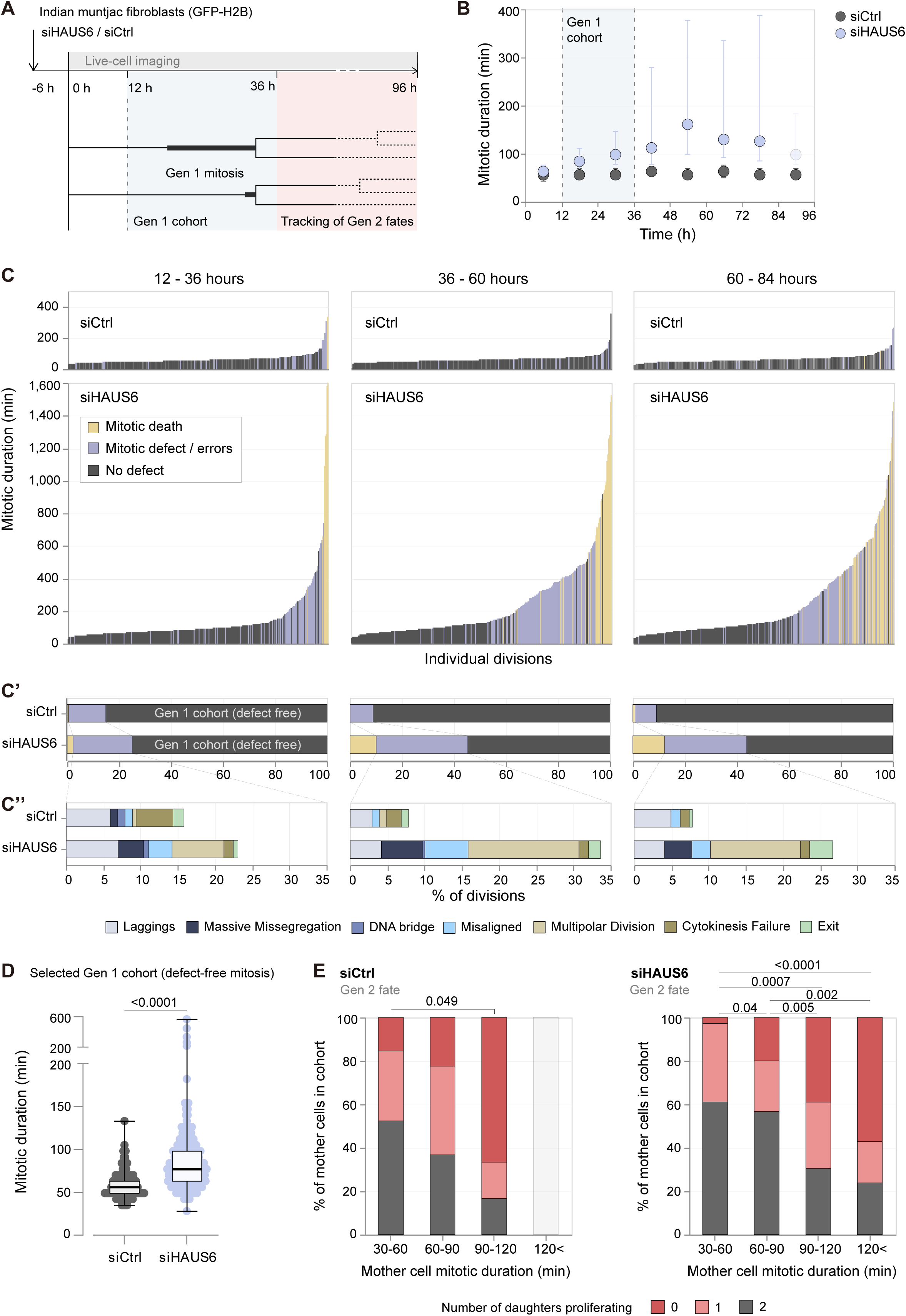
Delayed SAC silencing due to partial microtubule occupancy at kinetochores impacts daughter cell proliferation (A) Schematic summary of the live cell imaging setup and downstream analysis upon control or HAUS6 siRNA treatment. (B) Mitotic duration of Indian muntjac cells expressing H2B-GFP with decreasing HAUS6 levels (time of recording in the presence of control or HAUS6 siRNA; grey and blue, respectively). Data pooled from five independent experiments (siCtrl n = 823, siHAUS6 n = 1134). Circles and error bars denote median and interquartile range in each time bin (bin size = 12h). Note that after 84h of imaging mitotic durations are not accurate, as many cells remain in mitosis when the movie ends. (C) Mitotic cell fate as a function of mitotic duration in Indian muntjac cells treated with control or HAUS6 siRNA grouped into 24 h time bins. Bars show individual cells (siCtrl 12-36 h n = 202 cells, 36-60 h n = 204 cells, 60-84 h n = 243 cells, siHAUS6 12-36 h n = 316 cells, 36-60 h n = 309 cells, 60-84 h n = 322 cells, pooled from five independent experiments). (C’) Bars show the percentage of mitoses in C that divide error-free, with errors or die in mitosis. (C’’) Categories of mitotic defects found in C. Bars show percentage of total analyzed mitoses. (D) Data from C filtered for cells that divide without defects between 12 h to 36 h of recording (Gen 1 cohort). Dots show mitotic duration of individual cells (siCtrl n = 140 cells, siHAUS6 = 207, pooled from five independent experiments). Boxes show interquartile ranges, horizontal lines show median and error bars show min max. p <0.0001, Mann-Whitney test (two-sided). (E) Bars show the percentage of mother cells from the selected cohort shown in D that has 0, 1 or 2 daughter cells dividing at least once, as a function of mother cells time in mitosis in siCtrl (left) or siHAUS6 (right) treated populations. Data are binned (siCtrl 30-60 min n = 84 cells, 60-90 min n = 49 cells, 90-120 min n = 6 cells, >120 min n = 1 cell, siHAUS6 30-60 min n = 36 cells, 60-90 min n = 90 cells, 90-120 min n = 59 cells, >120 min n = 21 cells, pooled from five independent experiments). siCtrl 30-60 min vs 90-120 min, p = 0.05; siHAUS6 30-60 min vs 60-90 min, p = 0.04; 30-60 min vs 90-120 min, p = 0.0007; 30-60 min vs >120 min, p <0.0001; 60-90 min vs 90-120 min, p = 0.007; 60-90 min vs >120 min, p = 0.004, chi-square test with Benjamini-Hochberg multiple comparison correction. For all other comparisons p >0.05.

To determine the fate of daughter cells from mothers that *did* divide despite immature K-fibers, we defined a cohort of cells that underwent mitosis within 12 and 36 hours of recording and followed their offspring for 60 hours (Figure 6A). Since prolonged mitotic duration under low microtubule occupancy was tightly associated with mitotic errors (Figures 6C and 6C’) that may independently influence daughter cell fate ^66–69^, we sought to disentangle the two events. To this end, the low chromosome number of Indian muntjac fibroblasts allowed us to unequivocally exclude cells dividing with errors from our cell fate analysis (Figure 6D, 6E and 6E’). Importantly, although the remaining cohort only underwent a short Augmin depletion period (max. 42 hours) and divided without errors, siHAUS6-treated cells showed significantly prolonged mitotic times, presumably due to persistent low-microtubule occupancy (Figure 6D). Of note, within the control population, the progeny of cells that sporadically took longer to divide was also more likely to arrest proliferation then of cells that divided within the expected duration (15% [13/84] of cells with 30-60 min mitoses vs 67% [4/6] with 90-120 min mitoses have both daughters arresting; Figure 6E). However, the frequency of these events was significantly higher upon Augmin depletion (Figure 6D) where mother cells with increasing mitotic duration were increasingly more likely to give rise to daughter cells that stopped proliferating (3% [1/36] of cells with 30-60 min mitoses vs 39% [23/59] of cells with 90-120 min mitoses have both daughters arresting; Figure 6E). Consistently, we observed a similar gradual trend in cells released from up to 6 hours of mitotic arrest mediated by the Eg5/kinesin-5-inhibitor monastrol ^70^ (Fig. S7). Here, mother cells that experienced mitotic times longer than 100 minutes gave rise to daughters that showed increased cell cycle lengths or stopped proliferating (Fig. S7B), even following defect-free mitoses (5% [1/22] of cells with 30-60 min mitoses vs 35% [16/46] of cells with >120 min mitoses have both daughters arresting; Fig. S7E). Together, these results hinted at a conserved “mitotic stopwatch” surveillance of mitotic duration, in line with previous findings in human cells^29,31,32,34^.

Such a “memory” of mitotic duration was shown to result from the formation of a protein complex containing 53BP1, USP28 and p53 ^31,32,34^. To test whether this is conserved in Indian muntjac cells, we probed for mitotic stopwatch complex formation in cells arrested for up to 6h in the presence of monastrol (Figure 7A). Indeed, we were able to detect a significant accumulation of p53 and USP28 bound to 53BP1 in the mitotic population, but not in cells that remained adherent after drug treatment (LO population; Figure 7B), suggesting that the interaction specifically resulted from increased time in mitosis.

**Figure 7.**
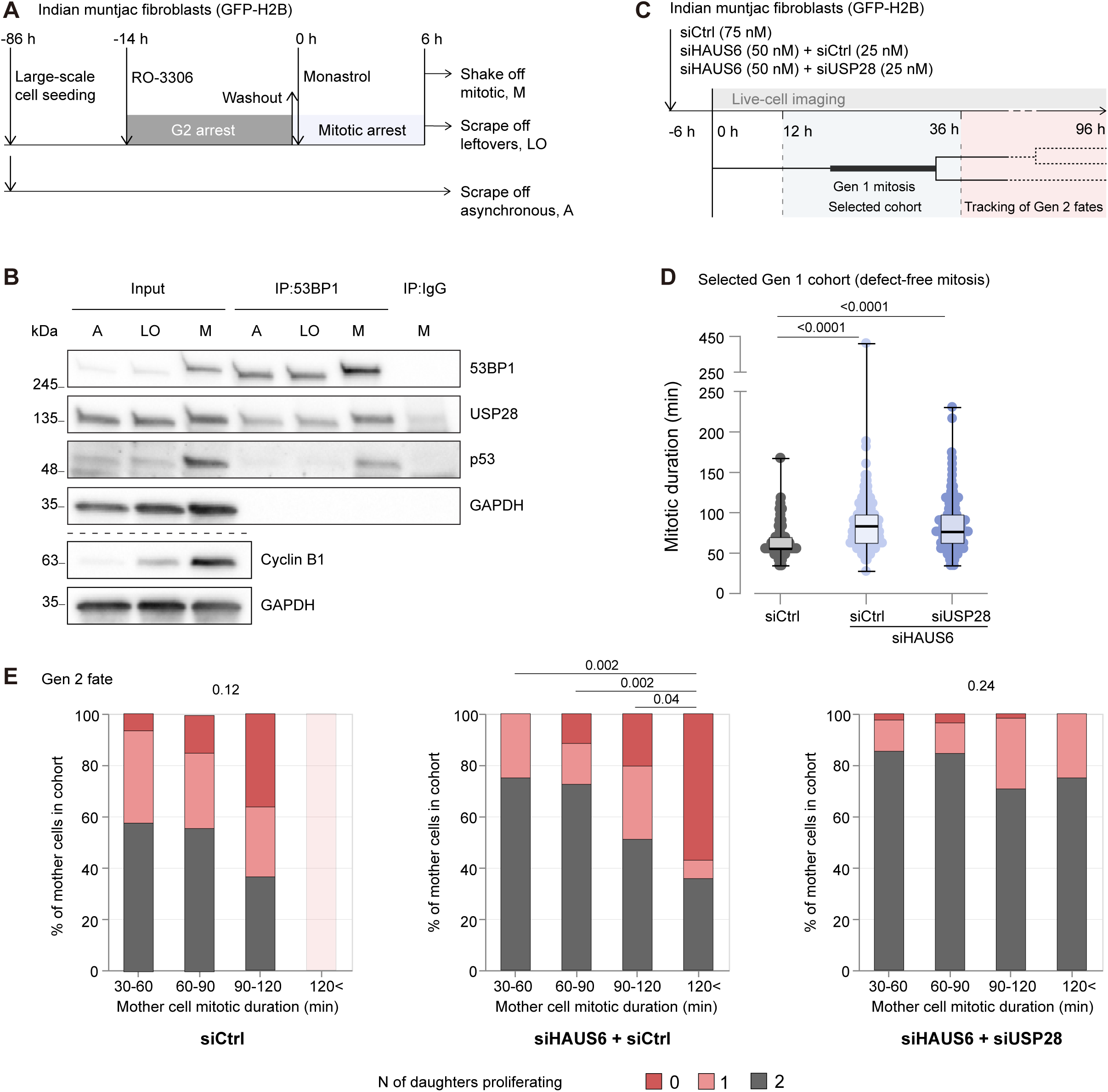
A conserved stopwatch mechanism stops proliferation of cells that delay mitotic exit due to low-microtubule occupancy at kinetochores (A) Schematic summary of co-immunoprecipitation assays after G2 synchronization and mitotic arrest with the indicated treatments. (B) Analysis of 53BP1 or IgG control immunoprecipitates (IP) in asynchronous (“A”), leftovers (“LO”) and 6 h mitotic arrested (“M”) Indian muntjac cells expressing H2B-GFP. Inputs correspond to whole cell lysates. GAPDH as loading control. Representative immunoblot from two independent experiments. (C) Schematic summary of the live cell imaging setup and downstream analysis upon HAUS6 in combination with control or USP28 siRNA treatment. (D) Mitotic duration of Indian muntjac cells expressing H2B-GFP treated with HAUS6 in combination with control or USP28 siRNAs. Dots show mitotic duration of individual cells from a select cohort - divide without defects between 12 h to 36 h of recording (Gen 1 cohort, siCtrl n = 96 cells, siHAUS6 + siCtrl = 153, siHAUS6 + siUSP28 n = 199, pooled from three independent experiments). Boxes show interquartile ranges, horizontal lines show median and error bars show min max. siCtrl vs siHAUS6 + siCtrl, p <0.0001; siCtrl vs siHAUS6 + siUSP28, p <0.0001, Kruskal-Wallis test with Dunn’s multiple comparisons test. For all other comparisons p >0.05. (E) Bars show the percentage of mother cells from the selected cohort that has 0, 1 or 2 daughter cells dividing at least once, as a function of mother cells time in mitosis in siCtrl (left), siHAUS6 + siCtrl (middle) or siHAUS6 + siUSP28 (right) treated populations. Data are binned (siCtrl 30-60 min n = 50 cells, 60-90 min n = 34 cells, 90-120 min n = 11 cells, >120 min n = 1 cell, siHAUS6 + siCtrl 30-60 min n = 20 cells, 60-90 min n = 69 cells, 90-120 min n = 49 cells, >120 min n = 14 cells, siHAUS6 + siUSP28 30-60 min n = 41 cells, 60-90 min n = 84 cells, 90-120 min n = 58 cells, >120 min n = 16 cells, pooled from three independent experiments). siCtrl p = 0.12, chi-square test; siHAUS6 + siCtrl 30-60 min vs >120 min, p = 0.002; 60-90 min vs >120 min, p = 0.002; 90-120 min vs >120 min, p= 0.04, for all other comparisons p >0.05, chi-square test with Benjamini-Hochberg multiple comparison correction; siHAUS6 + siUSP28 p = 0.24, chi-square test.

To test whether prolonged mitoses due to low microtubule occupancy at kinetochores were indeed registered and bookmarked by mitotic stopwatch complex formation, we co-depleted USP28 together with HAUS6 and again determined daughter fates (Figure 7C). Importantly, Augmin depleted cells took longer to divide then controls, irrespective of USP28 presence or absence (Figure 7D). Consistent with reports on USP28-deficient RPE-1 cells that were released from prolonged mitotic arrest ^71^, we found that siUSP28-treated cells lost stopwatch sensitivity and continued proliferating even after considerable mitotic arrest due to HAUS6 deficiency (2% [1/41] of cells with 30-60 min mitoses vs 2% [1/58] of cells with 90-120 min mitoses have both daughters arresting; Figure 7E). In conclusion, combining long-term cell fate analysis and biochemistry allowed us to pinpoint a conserved mitotic stopwatch mechanism in Indian muntjac cells, controlling daughter proliferation when SAC silencing is delayed due to low microtubule occupancy at kinetochores.

## Discussion

While there is a large consensus that SAC silencing can occur with low microtubule occupancy at kinetochores ^23–27,46^, whether and how SAC signaling along vertebrate kinetochores responds to highly localized microtubule attachments has remained a central open question. Current models suggest that mammalian kinetochores respond as a single unit that globally halts SAC signaling upon partial microtubule attachments ^25,26,46^. As so, low microtubule occupancy would cause kinetochores to respond as a whole and predicts a uniform loss of MAD1 along the kinetochores. Moreover, these switch-like models imply the existence of a sensitive microtubule “counting” mechanism that determines SAC silencing when a given occupancy at each individual kinetochore is reached. However, in virtue of the sub-diffraction nature of most vertebrate (humans included) kinetochores, this prediction had never been directly tested and the essence of a putative microtubule “counting” mechanism remains elusive. In this regard, Indian muntjac kinetochores offer a unique setting to test these ideas. In stark contrast with the predictions from switch-like SAC-silencing models, we found that the removal of the SAC protein MAD1 from Indian muntjac kinetochores is gradual, non-uniform, correlates with local microtubule density and kinetochore size, and depends on Augmin-mediated maturation of K-fibers.

In line with previous work ^46^, MAD1 signal decay from Indian muntjac kinetochores was well fit by a single exponential, indicating that SAC silencing in this system faithfully replicates established kinetics in other vertebrate systems. Switch-like models that favor independent microtubule binding at kinetochores and where a given percentage of occupancy is key to uniformly silence the SAC ^25,26^, predict that the fraction of unattached sites is expected to decay exponentially, with a timescale that does not depend on kinetochore size. Interestingly, however, we found that MAD1 decay on large kinetochores was 38% slower compared to small kinetochores. While the underlying reasons for these differences remain unclear, this finding suggests that switch-like models do not fully account for how the SAC is normally silenced. Indeed, both live-cell imaging and super-resolution CH-STED analysis in Indian muntjac cells revealed a non-uniform distribution of MAD1 on partially attached kinetochores that negatively correlated with microtubule occupancy. Moreover, laser microsurgery experiments demonstrated that once silenced, the SAC remains sensitive to highly-localized microtubule detachments from kinetochores, even when more than 50% occupancy persists. Lastly, experimental perturbation of microtubule occupancy at kinetochores by interfering with K-fiber maturation significantly delayed MAD1 removal and SAC silencing. Therefore, although low microtubule occupancy at kinetochores may eventually silence the SAC, there is an unavoidable impact on the respective mitotic duration. This has important implications in light of the recently-uncovered “mitotic stopwatch” mechanism, where even mild mitotic delays elicit a subsequent USP28-53BP1-p53-dependent cell cycle arrest in G1 ^29,31,32,35,72^, and the observation that SAC silencing with low microtubule occupancy increases the incidence of lagging chromosomes during anaphase, potentially leading to aneuploidy ^27^. As so, the mechanism underlying SAC silencing at individual kinetochores is likely to have evolved to maximize microtubule occupancy within a strict temporal window to avoid “bad memories” of mitosis, and we show here that the Augmin complex, through its role in K-fiber maturation ^43,65^, is instrumental in this process (see graphical abstract).

But how do partially attached kinetochores that are locally sensitive to microtubule attachments ultimately remove all MAD1/MAD2? In line with a model in which the number of unattached kinetochores ultimately determines the rate of MCC production that proportionally inhibits the APC/C ^8,9^, partially-attached kinetochores may produce less MCC than fully unattached kinetochores ^24^. Consequently, the APC/C would become sufficiently active and allow for slower, yet steady, Cyclin B1 degradation (and CDK1 inactivation), ultimately silencing the SAC after an extensive mitotic delay ^24^. Whether kinetochores eventually reach normal microtubule occupancy by the time the SAC is silenced after a delay remains unclear (see graphical abstract), but it should be noted that even cells that are unable to silence the SAC after extremely prolonged mitotic delays due to the complete absence of microtubules degrade Cyclin B1 and eventually exit mitosis due to residual APC/C activity ^73,74^. Our finding that CDK1 inhibition triggers SAC silencing irrespective of microtubule occupancy further supports this interpretation (see graphical abstract).

One may argue that the compound architecture of Indian muntjac kinetochores does not reflect what is normally found in other vertebrate species, including humans. However, we have no evidence that kinetochore architecture in Indian muntjac reflects a significant deviation at the level of the core microtubule-binding moduli, or other functional differences, including MAD1 decay during SAC silencing, when compared with other vertebrates ^41,43,46,75^. Noteworthy, recent works have revealed that many human, mouse and chicken centromeres/kinetochores also have a compound, often bipartite, architecture ^59,61,76,77^. In particular, the same human kinetochore could be found with one sub-domain end-on attached, and the other laterally attached to microtubules via the fibrous corona^76^. This is consistent with our finding of non-uniform, highly localized, MAD1 removal in response to local microtubule occupancy in Indian muntjac kinetochores. Importantly, while in these cases MAD1 and local microtubule density often negatively correlated along two apparently insulated adjacent domains on the same large kinetochore of chromosome 3+X (simply because these extreme cases are easier to detect), there were other cases (particularly evident after Augmin perturbation) where this negative correlation was observed along three or more sub-domains, without following a strict spatial pattern (also evident in the distinct responses after laser microsurgery experiments). Moreover, our finding that MAD1 is immobile within unattached kinetochores indicates that it can only be removed upon the establishment of highly-localized microtubule attachments (or eventually by a microtubule-independent SAC-silencing mechanism, such as MPS1 or CDK1 inactivation). Lastly, in line with our identification of the Augmin complex as a key determinant of microtubule occupancy at kinetochores linking timely SAC silencing with mitotic stopwatch surveillance, a recent CRISPR-Cas9 screen in human RPE1 cells independently uncovered the Augmin complex as an important factor controlling mitotic duration, perturbation of which triggered a mitotic stopwatch response ^78^. Overall, these results hint at functional conservation of the underlying molecular network linking SAC silencing with mitotic memory. Taken together, our data favors a model where anaphase onset is determined by the gradual silencing of the SAC in response to highly localized microtubule attachments along the entire kinetochore length, rather than a uniform, switch-like mechanism. This model reconciles gradual SAC silencing with increasing microtubule occupancy at kinetochores, without the need for a dedicated microtubule “counting” system that triggers SAC inactivation. Most relevant, our finding that microtubule occupancy at kinetochores is at the root between timely SAC silencing and mitotic memory, identifies a critical rate-limiting step with important implications for the control of cell proliferation in mammals.

## Materials and Methods

### Cell lines and culture conditions

Indian muntjac cell lines were grown in Minimum Essential Media (MEM) (Corning), supplemented with 10% FBS (GIBCO, Life Technologies). Indian muntjac hTERT-immortalized fibroblasts were a gift from Jerry W. Shay ^79^. IM fibroblasts stably expressing Venus-MAD1 were generated by lipofection of pVenus-MAD1 (kind gift from Jakob Nilsson ^80^). RPE-1 hTERT-immortalized cells were cultured in DMEM (Corning, 15323531) supplemented with 10% FBS (GIBCO, Life Technologies). All cells were kept at 37°C in humidified conditions with 5% CO_2_. Transfection was performed by incubating the cells with 1:400 Lipofectamine2000 (Invitrogen) in Opti-MEM (GIBCO, Life Technologies) for 6h, and stable and uniformly expressing cells were selected by FACS sorting (as detailed in ^42^. For the production of Venus-MAD1 and mScarlet-CENP-A co-expressing cell line, Venus-MAD1 cells were transduced with pLVx-mScarlet-CENP-A lentiviral plasmid (generated in ^43^). IM cells co-expressing 2x-GFP-CENP-A, GFP-CENTRIN-1 and mScarlet-MAD1 were produced by lentiviral transduction of pRRL-2x-EGFP-CENP-A (generated in ^43^), pLVX-EGFP-C1-CENTRIN-1 (kind gift from Manuel Thery; Addgene plasmid # 73331) and pInducer-mScarlet-MAD1 (generated in this paper). Cells expressing GFP-H2B were produced by lentiviral transduction of LV-GFP (kind gift from Elaine Fuchs; Addgene plasmid # 25999). Lentiviral transduction was performed as detailed in^42^. Cells were incubated with lentivirus particles in the presence of 1:2000 Polybrene (Sigma-Aldrich; TR-1003) for 24h before supplied with fresh complete media. Stable lines with sufficient fluorescence intensity were selected by FACS sorting.

### Plasmid design

pInducer-mScarlet-MAD1 was produced by cloning mScarlet (from pRRL-mScarlet-CENP-A) and MAD1 (from pVenus-MAD1) into the pInducer-20 backbone under a Tet-inducible promotor. To construct the empty backbone KASH-mCherry was removed by SalI + XhoI restriction digestion of pinducer 20 DN-KASH (kind gift from Daniel Conway; (Addgene plasmid # 125554)). Restriction sites were amplified from the plasmid by PCR (fwd: 5’-TTAAAGGAACCAATTCAGGCTAGCACGCGTATATCTAGACCCAGCTTTCT-3’; rev: 5’-TAAGCGTAGTCTGGGACG-3’) and added to the emptied vector through Gibson assembly. mScarlet and MAD1 were cloned into the NheI linearized pInducer-20 backbone through Gibson assembly.

### siRNA Experiments

Protein knockdown was performed as detailed in ^42^. Briefly, IM fibroblasts seeded at 70% confluency were starved with MEM supplemented with 5% FBS for 30 minutes before transfection. siRNA transfection was performed by adding Lipofectamine RNAi Max (1:400; Invitrogen) and 75 nM, 50 nM or 25 nM of siRNA in serum free-medium (Opti-MEM, Gibco) for 6h before replacing the solution with fresh complete media. Cells were analyzed 72 hours after depletion, unless specified otherwise. For the depletion of HAUS6, the following sequence was used: 5’-GGUUGGUCCUAAGUUUAUU[dT][dT]-3’^39^ at 50 nM. For the depletion of USP28, the following sequence was used: 5’-GCTTCCGGACATGTTGAAATA[dT][dT]-3’ at 25 nM. Cells mock transfected (lipofectamine only) or transfected with luciferase-targeting siRNA (5’-CAUUCUAUCCUCUAGAGGA[dT][dT]-3’) at 75 nM, 50 nM or 25 nM were used as controls.

### Drug Treatments

For acute inhibition of Mps1 and CDK1 kinases, 4 µM MPS1-IN-1 (kind gift from N.Gray ^51^) and 10 μM RO 3306 (ChemCruz; CAS 872573-93-8), respectively, were added shortly after Nuclear Envelop Breakdown (NEB). To arrest cells in G2, these were treated with 10 μM RO 3306 for 14h. Microtubule depolymerization was triggered using 1 µM of Nocodazole (Sigma-Aldrich; CAS 31430-18-9) for 30 mins before the analysis. To increase mitotic duration cells were arrested in prometaphase with 75 or 100 μL of the EG5 inhibitor, Monastrol (Tocris Bioscience, Cat. No.1305) for 6h. Metaphase arrest was obtained using 5 µM MG132 (Sigma-Aldrich; CAS 133407-82-6). Fixed cell analysis using MG132 was performed in the first 1.5 h after drug addition to avoid cohesion fatigue. SiR-tubulin (Spirochrome; SC002) ^81^ was used to visualize microtubules, at 50 nM concentration incubated for 1 h prior to live-cell imaging.

### Immunofluorescence

Indian muntjac fibroblasts were seeded on fibronectin coated coverslips (#1.5 thickness) 24h before the experiment. Depending on the microscopy routine, different fixation protocols were used: for widefield imaging cells were incubated with 4% paraformaldehyde (Electron Microscopy Sciences) in PBS for 10 min at room temperature; for STED microscopy PFA 4% supplemented with 0.1%-0.2 % Glutaraldehyde (Electron Microscopy Sciences) in Cytoskeleten Buffer (CB - 274 mM NaCl, 10 mM KCl, 2.2 mM Na_2_HPO_4_, 0.8 mM KH_2_PO_4_, 4 mM EGTA, 4 mM MgCl_2_, 10 mM Pipes, 10 mM glucose, pH 6.1) for 10 min at room temperature was used; for same cell correlative live confocal and super-resolution STED microscopy pre-warmed 2x concentrated PFA + Glutaraldehyde diluted in imaging media to a final concentration of 4% and 0.2%, respectively, was added to the imaging chamber. After a 5 min incubation at the microscope (37°C), the fixation solution was replaced with PFA 4% supplemented with 0.2 % Glutaraldehyde in CB for an additional 5 min at room temperature. Autofluorescence was quenched by a 0.1% sodium borohydride solution (Sigma-Aldrich) for 7min. Cells were permeabilized with CB-0.5%Triton for 20-30 min and blocked with CB-0.05% Tween 20 with 10% FBS (blocking buffer) for 1 hour at RT. Samples were incubated with primary antibodies diluted in blocking buffer over-night at 4°C. The following primary antibodies were used: human anti-centromere antiserum (ACA, Fitzgerald; 90C-CS1058) 1:2000/1:200 (Widefield/ STED); anti-tyrosinated tubulin (Bio-Rad; MCA77G) 1:2000/1:100 (Widefield/ STED); anti α-tubulin (clone B-5-1-2, Sigma-Aldrich; T5168) 1:100 (correlative live-cell and fixed STED microscopy). Cells were washed with PBS-0.05% Tween before incubation with the corresponding secondary antibodies for 1h at RT – Alexa Fluor 568 and 647 (Thermo Fisher Scientific) 1:1000; or abberior STAR 580 (Abberior Instruments; ST580) and abberior STAR-Red (Abberior Instruments; STRED) 1:200 for STED microscopy. DNA was labeled with a quick incubation in the presence of 1 µg/mL 4’6’-Diamidino-2-phenylindole (DAPI) in PBS-0.05% Tween. The samples were washed in PBS and mounted on glass slides with a mounting solution (20 mM Tris pH8, 0.5 N-propyl gallate, 90% glycerol).

### Monitoring kinetochore MAD1 levels in live recordings

Indian muntjac fibroblasts stably expressing Venus-MAD1 or co-expressing Venus-MAD1 and mScarlet-CENP-A were plated on fibronectin coated glass bottom 35 mm FluoroDish (World Precision Instruments; FD35-100), 36-48h before imaging. Before imaging, normal culture media was replaced with Leibovitz’s L15 medium (GIBCO, Life Technologies) supplemented with 10% FBS and the cell-permeable microtubule dye, SiR-tubulin. Live-cell imaging was performed on a temperature-controlled Nikon-Ti microscope equipped at the camera port with a Yokogawa CSU-X1 spinning-disc head with Borealis and an iXon+ DU-897 EM-CCD (Andor). Cells were imaged with an oil-immersion 60x 1.4 NA Plan-Apo DIC CFI objective (Nikon, lambda series), yielding a 176 nm/pixel sampling, or a 100x 1.4 NA Plan-Apo DIC CFI objective (Nikon, VC series), yielding a 106 nm/pixel. Images were acquired using 488 nm, 561nm and 647 nm (Coherent) laser lines and acquisition was controlled by NIS Elements AR software. To ensure that kinetochore could be tracked overtime with a reduced number of optical sections, cells were imaged under modest cell confinement. An image stack was acquired: every 6 seconds, 9 planes separated by 0.5 µm (Figures 1, 6 and Fig. S1C); 1 min, 11 planes separated by 0.5 µm, Fig. S1A, B); 10 seconds, 9 planes separated by 0.5 µm (Figure 3). Fluorescence intensity of kinetochore Venus-MAD1 was manually tracked throughout mitosis using the image analysis software Fiji 2.16.0 ^82^. The recorded image stacks were sum projected and a region was drawn encircling the kinetochore and three different cytoplasmic background sites at each timepoint. MAD1 kinetochore fluorescence intensity (S_KT_) was calculated by subtracting the averaged cytoplasmatic signal (S_BG_), represented in the following equation: S_KT, corrected_ = S_KT_ - S_BG, averaged_×A_KT_/A_BG_ S: raw integrated density; A: Area. The calculated signal intensity was normalized to the followings: the clear last peak of the signal decay curve in untreated/control, mock treated and siHAUS6 conditions; the first frame after adding the inhibitor in MPS1i and CDK1i treatments; the average of the first 10 timepoints in nocodazole treated cells. A single-phase exponential decay function was fitted globally across curves polled from all kinetochores in each experimental condition using the equation: Y = (Y0-Plateau) x (exp^(-Kx)) + Plateau, in GraphPad Prism 9 (Boston, Massachusetts USA). The half decay time (t_1/2_) for each condition was derived from the fit according to t_1/2_ = ln(2)/K. To test for differences in the rate of MAD1 decay (K, derived from the global fit) in untreated vs treated cells or large vs small kinetochores between conditions an extra sum-of-squares F-test was used. Quantification of the signal intensity ratio between kinetochore and cytoplasm (Fig. S1B), was performed manually using Fiji 2.16.0. Three regions were drawn encircling the kinetochore (KT), the cytoplasm (cyto) and an area outside of the cell (background signal, BG) at define time points (anaphase onset, 10, 20, 30 and 40 minutes before anaphase). The kinetochore and cytoplasmic fluorescence intensities were calculated by subtracting background signal, represented in the following equations: S_KT, corrected_ = S_KT_/A_KT_ - S_BG_/A_BG_; S_cyto, corrected_ = S_cyto_/A_cyto_ - S_BG_/A_BG;_ S: signal; A: Area. The ratio of the corrected kinetochore over cytoplasmatic signals was then calculated. To observe how Mad1 signal spatial distribution within the kinetochore changed throughout time kymographs were produced from live recordings of IM cells (Figure 3). Image stacks were sum projected using Fiji 2.16.0 and kymographs were generated with a custom-written routine in MATLAB (Mathworks, Natick, MA) that compensates for spindle rotation and translations previously described in ^64^.

### Cell confinement

For cell confinement, we adopt a cell confiner as previously described ^39, 62^ using a custom-designed polydimethylsiloxane (PDMS, RTV615, GE) layout to fit a 35mm diameter fluorodish. A suction cup was custom-made with a 10:1 mixture (w/w PDMS crosslinker A/ B) and baked on an 80 °C hot plate for 30mins and left to dry over-night before unmolding. A PDMS confinement slide was then prepared with a 10:1 mixture (w/w PDMS crosslinker A/ B) molded into micropillars with a height of 8 μm (using a SU-8 wafer mold) polymerized onto a 10 mm round coverslip and baked for 95 °C for 15 min. The PDMS confinement slide was attached to the PDMS suction cup and connected to a vacuum generator (AF1-dual, Elveflow).

### Quantification of MAD1 intra-kinetochore mobility via fluorescence recovery after photobleaching (FRAP)

For FRAP experiments, IM cells stably expressing Venus-MAD1 were plated on fibronectin coated 35mm FluoroDish (World Precision Instruments, FD35-100) 36-48 h before imaging. To elicit uniform and maximal kineotchore-MAD1 occupancy, cells were treated with nocodazole for 30 minutes before the experiment. To ensure that the bleached kinetochore could be tracked overtime with a reduced number of optical sections, an agar overlay was placed on top of the cells decreasing the cell volume, as described in ^63^. Briefly, a 170 μm thick layer of 2% low-melting-point agarose in L15 medium supplemented with 10% FBS was prepared in advance and incubated for 30 minutes with pre-warmed L15 + 10% FBS containing nocodazole. Immediately before imaging, a small piece of the agar layer was gently placed over the cells that were maintained in a minimal amount of medium to prevent cell dehydration while ensuring compression. FRAP experiments were conducted on a Leica Scanning Confocal STELLARIS 8 FALCON (Leica Microsystems, Germany), equipped with an incubation chamber and a power HyD S detector. Cells were imaged with a water-immersion 86x 1.20 NA HC PL APO STED white objective (Leica Microsystems, Germany), plus a 4.5x zoom, yielding an 86 nm/pixel sampling. The 488-nm emission wavelength of a white Light Laser (WLL) was used to bleach MAD1 signal from either part of or an entire kinetochore with 2 pulses of 0.3 seconds each. An image stack of 3 z-planes at a 0.5 μm interval was acquired with the following routine: 3 pre-bleach frames, every 1.3 seconds; kinetochore bleaching; post-bleach 5 frames every 1.3 seconds, 10 frames every 2 seconds and then every 3 seconds for up to 55 seconds. The selected representative images show sum-intensity projections of the image stacks. Venus-MAD1 fluorescence at the kinetochore was manually tracked on the sum projected image stacks using Fiji 2.16.0and a custom-designed script in Python 3.13. Four rectangular ROIS were defined corresponding to: (i) the bleached region of the experiment kinetochore; (ii) the non-bleached region of the experiment kinetochore; (iii) the unperturbed kinetochore from the pair (sister KT) and (iv) a kinetochore from an unperturbed pair (control KT). Additionally, two square ROIs with 0.86 μm sides were drawn over the cytoplasm to measure background fluorescence (BG). A custom-designed script was used to draw the kinetochore ROIs with a constant width of 0.6 μm, and length defined by the distance of the kinetochore extremities in the pre-bleach frame (manually marked based on MAD1 signal using Fiji 2.16.0). In the case of the partially ablated kinetochore the length of the non-bleached region was defined at the first frame post-bleaching, and the length of the bleached region was defined as the difference between the total kinetochore length and the length of the non-bleached region. The long axis of the rectangle was defined based on the vector described by the points marking the extremities of the non-bleached signal and was adjusted at every time point to track the kinetochore motion. MAD1 kinetochore fluorescence intensity (S_KT, corrected_) was corrected for fluctuations in the cytoplasmatic fluorescent signal as described in the following equation: S_KT, corrected_ = S_KT_ / (a x time + b); S_KT_: kinetochore MAD1 raw integrated density; a, b: constants provided by a linear fit of the cytoplasmatic MAD1 raw integrated density. The corrected kinetochore signal was then normalized to the average of the 3 pre-bleach values (S_KT, normalized_) and fit to a single exponential association function: Y = Y0 + (Plateau – Y0) x (1-(exp^(-Kx))); Y0 value was constrained to be the S_KT, corrected_ at the first frame post-bleaching. Half recovery time (t_1/2_) was derived from the fit according to t_1/2_ = ln(2)/K. Fluorescence recovery percentage (Recovery) was calculated according to Recovery = (Plateau – Y0) / (1 – Y0). Fitting and downstream statistical analysis was performed using GraphPad Prism 9 (Boston, Massachusetts USA). Cells where bleaching efficiency was < 50%; focal plane changes were > 1 μm; or where the normalized MAD1 fluorescence could not be fitted to a single exponential (if the 95 % confidence interval for any of the parameters could not be determined) were excluded from the analysis. Line intensity profiles showing Venus-MAD1 signal along the kinetochore were measured from a ROI covering the entire kinetochore length and a width of 0.250 μm at selected times.

### Quantification of kinetochore MAD1 signal distribution with Stimulated Emission Depletion (STED) Microscopy

Indian muntjac fibroblasts stably expressing Venus-MAD1 were seeded on glass coverslips and processed as described in the “Immunofluorescence” section. 3D image stacks were acquired using an Abberior’Expert Line’ gated-STED microscope, equipped with a Nikon Lambda Plan-Apo 1.4NA 60x objective lens. CH-STED mode was implemented as described before ^62^. Images were acquired using excitation wavelengths at 488 nm, 561 nm, and 640 nm and a single doughnut-shape depletion beam at 775 nm, a 0.8 Airy unit pinhole and the STED channel had a time-gate threshold of 500 ps. Pixel size was set to 30 nm in xy and 200 nm in z and Image stacks covering near full spindle volume (∼8 μm). The selected representative images show sum-intensity projections of a fraction of the Z-slices (5-10). In images that show merged channels, histograms were cropped to allow visualization of all structures. Venus-MAD1 and tubulin signal distribution was analyzed along individual kinetochores in sum projected 3D stacks (5-10 slices) using Fiji 2.16.0. A line of 0.6 µm width was drawn along the longitudinal axis of the kinetochore (defined by the ACA signal). Intensity line profiles were extracted along the kinetochore for both MAD1 and tubulin channels and background subtracted (average cytoplasmatic intensity value). Corrected signal intensity was normalized to the maximum intensity value within each kinetochore. To visualize and measure correlations between MAD1 and tubulin signals, we calculated the intensity z-scores, standardized internally (for each kinetochore). Each position along the kinetochore is represented as a point in a 2D space where the z-scores are the x-y axes and an ensemble of kinetochores was either overlaid in the same scatter plot or displayed and calculated for isolated kinetochores. The product zMAD1.ztubulin at each data point is the local contribution to an overall correlation. Averaging over all data points yields the covariance normalized to the standard deviation product, known as the Pearson correlation coefficient. The Matlab function *corrcoef* was used, which calculates Pearson’s correlations and p-values with a two-tailed Student t-test for zero correlation. 3D reconstructions of the image stacks were obtained with Imaris 9.3.1 (Oxford Instruments).

### Laser microsurgery and correlative live-cell confocal and fixed-cell CH-STED microscopy

Indian muntjac cells expressing 2x-GFP-CENP-A and GFP-CENTRIN-1 and mScarlet-MAD1 were seeded on fibronectin coated 25 mm, no. 1.5, round coverslips (CAT 10593054, fisher scientific) 48h prior to the experiment. To induce mScarlet-MAD1 expression, cells were incubated with 1 ug/mL of doxycycline for 24h. Before imaging the coverslips were mounted on a 35 mm Attofluor Cell Chamber (CAT A7816, Invitrogen) and the media replaced with in Leibovitz’s L15 medium (GIBCO, Life Technologies) supplemented with 10% FBS. Cells were imaged using Nikon-Ti spinning-disk confocal microscope with a 100x 1.4 NA oil objective as detailed in the “Monitoring kinetochore MAD1 levels in live recordings” section. Cells in metaphase were identified by having MAD1 negative kinetochores aligned at the equator and increased inter-kinetochore and pole-pole distance (assessed by GFP-CENTRIN-1). After recording a pre-surgery image stack, one of the large kinetochores (from the IM chromosome 3+X) was partially ablated with a 532 nm laser, controlled by a custom routine. A detailed description of the microsurgery setup can be found in ^83^. 2-5 consecutive pulses with a 0.35 μm step and a 12 Hz repetition rate) were applied. The pulse width was 10 ns and the pulse energy was 3.9–4.4 μJ. After surgery an image stack (5 planes, 0.75 μm z-step) was recorded every minute until MAD1 signal was detected at the kinetochore, for up to 30 min; after MAD1 detection the acquisition routine was changed: 7 planes, 0.5 μm z-step every 20 seconds for 5 min. As a control for uniform MAD1 signal distribution, non-ablated cells were imaged upon acute nocodazole treatment (5 to 60 min incubation). For correlative live-cell confocal and fixed CH-STED microscopy analysis, once MAD1 signal was detected at the kinetochore a fixation solution was added to the imaging chamber. The location of the cell of interest was marked with a laser-engraved pattern on the glass coverslip. The sample was processed for staining as described in the “Immunofluorescence” section. After locating the surgery cell with a 10x objective, a super-resolved image stack of the spindle was acquired with CH-STED as described in the Quantification of kinetochore MAD1 signal distribution with Stimulated Emission Depletion (STED) Microscopy” section (pixel size 35 nm in xy and 150 nm in z). Representative images show a sum (live-cell data) or max-intensity (fixed-cell data) projection of all z-slices that contain the kinetochore of interest. Histograms were adjusted per frame to allow better visualization of the representative phenotypes. To measure MAD1 recruitment after laser microsurgery, kinetochores were manually tracked using the GFP-tagged CENP-A as a reference in Fiji 2.16.0. kinetochore deformation was defined as the first time point when an increase in the angle between the line connecting the ends of the surgery kinetochore and the line connecting the ends of the unperturbed kinetochore from the pair (sister kinetochore) was observed. MAD1 recruitment was defined as the first time point when mScarlet-MAD1 signal was visible at the sister kinetochore. MAD1 signal distribution along the sister kinetochore was measured at the time point when signal intensity appeared highest, based on visual assessment. On the cells where correlative live-cell spinning-disk confocal and CH-STED microscopy was performed, tubulin signal distribution was measured in the fixed sample. Measurements were performed on sum projected stacks containing the kinetochore pair of interest. A segmented line of 0.8 μm width (for MAD1, live imaging) or 0.9 μm width (for tubulin, fixed cell imaging) was drawn to span the full length of the kinetochore, as defined by the CENP-A signal, starting from the end that retained its counterpart (or randomly in nocodazole treated cells). Intensity line profiles were extracted along the kinetochore for all channels and background subtracted (cytoplasmatic signal near the kinetochore). The corrected intensity profile was normalized to the maximum within each kinetochore, as exemplified for CENP-A: S_CENP-A, norm_ = (S_CENP-A_-S_BG CENP-A_) / max (S_CENP-A_ - S_BG CENP-A_), S: mean pixel intensity. To correct for intensity variations caused by kinetochore deformation or movement, intensity profiles are shown as a ratio of MAD1/tubulin over CENP-A. Kinetochore length was normalized to a relative scale ranging from 0 (start of the line profile) to 1 (end of the line profile). The average MAD1/CENP-A ratio intensity profile for each condition (Nocodazole vs Surgery), was calculated using the ‘Multiple average curves’ function with linear interpolation on OriginPro 2022 (OriginLab Corporation, Northampton, MA, USA). The coefficient of variation of the MAD1/CENP-A ratio was calculated as the standard deviation of the MAD1/CENP-A intensity values along each kinetochore, divided by their mean. Cells that failed to recruit MAD1 within the mean recruitment time + 1 s.d. (∼12 minutes) or exhibited only very faint MAD1 signal were excluded from the MAD1 distribution analysis. Statistical analysis and plots were done using GraphPad Prism 9 (Boston, Massachusetts USA).

### Quantification of kinetochore MAD1 levels in prometaphase and metaphase HAUS6 depleted cells

Control and HAUS6 depleted cells expressing Venus-Mad1 were treated with MG 132 for 1h and then fixed and stained for as detailed in the “Immunofluorescence for wide-field” section. Samples were imaged on AxioImager Z1 equipped with a CCD camera (ORCA-R2, Hamamatsu) operated by Zen software (Carl Zeiss, Inc.). 3D image stacks covering the entire cell volume (0.23 μm z-step) were acquired using a 100× Plan-Apochromatic oil differential interference contrast objective lens, 1.46 NA (Carl Zeiss Microimaging Inc.) and excitation wavelengths at 488 nm, 561 nm, and 640 nm. The number of cells with MAD1 positive kinetochores was visually assessed for each condition. Kinetochore Venus-MAD1 signal intensity was measured on a single focal plane using Fiji 2.16.0. Cytoplasmatic Venus-MAD1 signal was averaged from 3 regions outside the spindle area. The ration between kinetochore and cytoplasm signal was calculated according to the equation: S_KT_ / S_cytoplasm;_ S: mean pixel intensity. Statistical analysis and plots were done using GraphPad Prism 9 (Boston, Massachusetts USA).

### Cell tracking and daughter fate analysis in live recordings

Indian muntjac fibroblasts stably expressing GFP-H2B were plated on fibronectin coated µ-Slide 4 Well Glass Bottom (Ibidi GmbH, 80427) with Ham’s F-10 media supplemented with 20% FBS 48h before imaging. Cells were recorded using the Zeiss LSM 980 Airyscan confocal microscope built on an Axio Observer 7 inverted stand with a motorized piezo stage and active focus stabilization. A controlled atmosphere chamber (CO2 and temperature) was used. Objective lens was a dry 0.3NA 10x Plan Neofluar. Detection was performed with two GaAsP detectors. Image acquisition was carried out with ZEN 3.11 software. 3D image stacks (pixel size 830 nm in xy, 4 planes separated by 6.7 μm, pinhole Airy unit of 1.23, scan speed of 7, no averaging) were recorded every 7 min using both transmitted light and an excitation wavelength at 488 nm. In experiments where mitotic delay was induced through monastrol treatment, cells were imaged in the presence of the drug for 6h before its wash out into fresh media. The fate of cell families was tracked for 72h. In experiments where USP28 and/or HAUS6 was depleted, cells were treated with control, USP28 or HAUS6 siRNAs 6h before imaging. Cell families were then tracked for 96h. Cells were tracked manually using TrackMate ^84^. Single-cell lineages and cell cycle durations (frame of anaphase mother cell to frame anaphase daughter cell) were analysed by modifying previously published ^85^ Jupyter notebooks (7.0.8). We defined mitotic duration as the frame of mitotic onset (condensing chromatin) until the first frame of anaphase (clearly separated daughter chromatin). For each division event, we determined mitotic defects and errors, and the fate of the daughters. For daughter fate analysis tracks where daughter cells migrated out of the field were excluded. For binned data, bins with n < 3 cells were greyed out or not shown. Data analysis, filtering and plotting was integrated using Jupyter notebooks (Python 3.12) and the pandas (2.2.2) ^86^ and Altair (5.4.1) ^87^ libraries. Fiji/ImageJ ^82^ macros.

### Co-immunoprecipitation

Indian muntjac fibroblasts stably expressing GFP-H2B were grown on with Ham’s F-10 media supplemented with 20% FBS. To obtain a mitotic enriched population, cells were synchronized in G2 with RO-3306 for 14h and then released into fresh media supplemented with monastrol for 6h. 10 million mitotic cells were collected by shake-off. The adherent population after shake-off (leftovers, LO) and untreated (asynchronous) cells were collected by cell scrapping. Cells were pelleted at 300 g for 5 minutes, washed with cold PBS and cell pellets were snap frozen in liquid nitrogen. A co-immunoprecipitation assay was optimized from the protocol described in ^29^. Cell pellets were resuspended in lysis buffer (20 mM Tris/HCl, pH 7.5; 50 mM NaCl; 0.5% Triton X-100; 5 mM EGTA; 1 mM dithiothreitol; 2 mM MgCl_2_; Pierce Protease and Phosphatase inhibitor mini tablets, EDTA-free [Thermo Fisher Scientific; A32961]) and incubated for 45 minutes at 4°C with agitation. Cell lysates were centrifuged at 15 000 rpm for 15 minutes at 4°C to collect the supernatant. Protein concentration was quantified and a lysate solution containing 1.5 mg of protein in 1 mL of lysis buffer was prepared for each condition. Lysates were incubated with 3 μg of anti-53BP1 antibody (Novus, NB100-304) or control rabbit IgG (Antibodies Incorporated, 43-630-0010) for 2h at 4°C with agitation and subsequently with 20 μL of Pierce Protein A Magnetic Beads (Thermo Fisher Scientific, 88845), pre-washed 3 times with lysis buffer, for 1h at 4°C with agitation. The non-binding supernatant faction was collected and the beads were washed 4 times with lysis buffer. The bound fraction was eluted from the beads with 30 μL of Laemmli sample buffer (100 mM Tris-HCl, pH 6.8, 4% SDS, 20% glycerol, 10% β-mercaptoethanol and 0.01% bromophenol blue) for 5 min. After separated from the beads, the eluate was denatured at 95 °C for 5 min and the samples were analyzed through western blot.

### Western blotting

Whole protein samples (input) containing 40 ug in Laemmli sample buffer (50 mM Tris-HCl, pH 6.8, 2% SDS, 10% glycerol, 5% β-mercaptoethanol and 0.005% bromophenol blue) or co-IP eluate samples were loaded on a 4–15% Mini-PROTEAN® TGX Precast Protein Gel (Bio-Rad, 4561084), mounted on a Mini-PROTEAN vertical electrophoresis apparatus (Bio-Rad). NZYColour Protein Marker II (NZYTech, MB09002) was used as a ruler. Blotting was performed with a Transfer-Blot Turbo transfer system (Bio-Rad) onto 0.2 µm PVDF membranes (Bio-Rad, 1704156). Membranes were blocked in blocking buffer (5% powder milk in PBS Tween 0.1%) for 1h. Anti-53BP1 mouse 1:1000 (Millipore, 05-725), anti-USP28 rabbit (Abcam, ab126604), anti-p53 mouse 1:100 (Thermo Fisher Scientific, MA1-12549), anti-Cyclin B1 mouse 1:1000 (Cell Signaling Technology, 4135) and anti-GAPDH mouse 1:10 000 (Proteintech, 60004-1) primary antibodies were diluted in blocking buffer and incubated 1 h at room temperature with agitation. Anti-rabbit HRP (Jackson ImmunoResearch, 111-035-003), anti-mouse HRP (Jackson ImmunoResearch, 115-005-003) and anti-mouse light chain specific HRP (Jackson ImmunoResearch, 115-035-174) secondary antibodies were diluted 1:5000 in blocking buffer and incubated 1 h room temperature with agitation. Membranes were developed with Clarity Western ECL Blotting Substrate (Bio-Rad, 1705061) and detected in the Chemidoc XRS system and Image Lab software (Bio-Rad).

## Resource availability

### Lead contact

Further information and requests for resources and reagents should be directed to and will be fulfilled by the lead contact, Helder Maiato.

### Materials availability

All reagents generated in this study are available from the Lead Contact without restriction.

### Data and code availability

Any additional information required to reanalyze the data reported in this paper is available from the lead contact upon request.

Analysis code is available as public repositories at https://github.com/CID-LAB-i3S/KinoFRAP and https://github.com/CID-LAB-i3S/MitoticMemory.

## Supplemental information

Document S1. Figures S1-S7

**Video S1**, related to Figure 1

MAD1 removal from kinetochores during an unperturbed mitosis (prometaphase to metaphase). Confocal recording of an Indian muntjac fibroblast expressing Venus-MAD1 (orange) and with SiR-tubulin labeled microtubules (grey). Scale bar: 5 µm. Time stamps are min:sec.

**Video S2**, related to Figure 1

MAD1 remains at kinetochores upon Nocodazole treatment. Confocal recording of an Indian muntjac fibroblast expressing Venus-MAD1 (orange) and with SiR-tubulin labeled microtubules (grey). Scale bar: 5 µm. Time stamps are min:sec.

**Video S3**, related to Figure 1

MAD1 removal from kinetochores in a prometaphase cell treated with MPS1 inhibitor. Confocal recording of an Indian muntjac fibroblast expressing Venus-MAD1 (orange) and with SiR-tubulin labeled microtubules (grey). Scale bar: 5 µm. Time stamps are min:sec.

**Video S4**, related to Figure 1

MAD1 removal from kinetochores in a prometaphase cell treated with CDK1 inhibitor. Confocal recording of Indian muntjac fibroblasts expressing Venus-MAD1 (orange) and with SiR-tubulin labeled microtubules (grey). Scale bar: 5 µm. Time stamps are min:sec.

**Video S5**, related to Figure 2

MAD1 fluorescence recovery after partial kinetochore bleaching. Confocal recording of an Indian muntjac cell pre-and post-bleach of part a kinetochore. Dashed line boxes indicate the ablated kinetochore (non-bleached region - dark blue; bleached region – light blue). Scale bar: 2 µm. Time stamps are min:sec.

**Video S6**, related to Figure 2

MAD1 fluorescence recovery after total kinetochore bleaching. Confocal recording of an Indian muntjac cell pre-and post-bleach of part an entire kinetochore. Dashed line box indicates the ablated kinetochore (yellow). Scale bar: 2 µm. Time stamps are min:sec.

**Video S7**, related to Figure 4

Laser microsurgery ablation results in localized kinetochore MAD1 recruitment. Confocal recording of an Indian muntjac cell expressing 2x-GFP-CENP-A and GFP-CENTRIN-1 (both in blue) and mScarlet-MAD1 (orange) upon partial kinetochore surgery. Arrow and asterisks points to the ablated kinetochore region pre-and post-surgery, respectively. Scale bar: 2 µm. Time stamps are min:sec.

**Video S8**, related to Figure 5

MAD1 removal from kinetochores during mitosis in a mock treated cell (prometaphase to metaphase). Confocal recording of Indian muntjac fibroblasts expressing Venus-MAD1 (orange) and with SiR-tubulin labeled microtubules (grey). Scale bar: 5 µm. Time stamps are min:sec.

**Video S9**, related to Figure 5

MAD1 removal from kinetochores during prometaphase in a siHAUS6 treated cell. Confocal recording of Indian muntjac fibroblasts expressing Venus-MAD1 (orange) and with SiR-tubulin labeled microtubules (grey). Scale bar: 5 µm.

Time stamps are min:sec.

## Acknowledgements

We would like to thank all colleagues that generously provided reagents used in this study, and Paula Sampaio and Maria Azevedo from i3S Advanced Light Microscopy Facility for technical support. We are indebted to Franz Meitinger for technical advice on the purification of the mitotic stopwatch complex. We also thank CID Lab members for discussions and constructive feedback throughout the course of this project, with special thanks to Daniela Andrade and Rui Martins for their strong contribution during mitotic shake-offs. NO, JS-O and EW were respectively supported by fellowships SFRH/BPD/118126/2016, SFRH/BD/07730/2021 and 2022.13008.BD from Fundação para a Ciência e a Tecnologia of Portugal. This project was funded by The European Union through the European Research Council Advanced grant KAREVO (project 101140624) to HM.

## Author contributions

Conceptualization: HM

Methodology: NO, JS-O, TK, AJP, EW

Investigation: NO, JS-O, TK

Visualization: NO, JS-O, TK, AJP

Funding acquisition: HM

Project administration: HM

Supervision: HM, AJP

Writing – original draft: NO, HM

Writing – review & editing: NO, JS-O, TK, AJP, EW, HM

