## Supplementary figures and legends for "Microtubule occupancy at kinetochores links checkpoint silencing with mitotic memory"

### Supplemental Figures

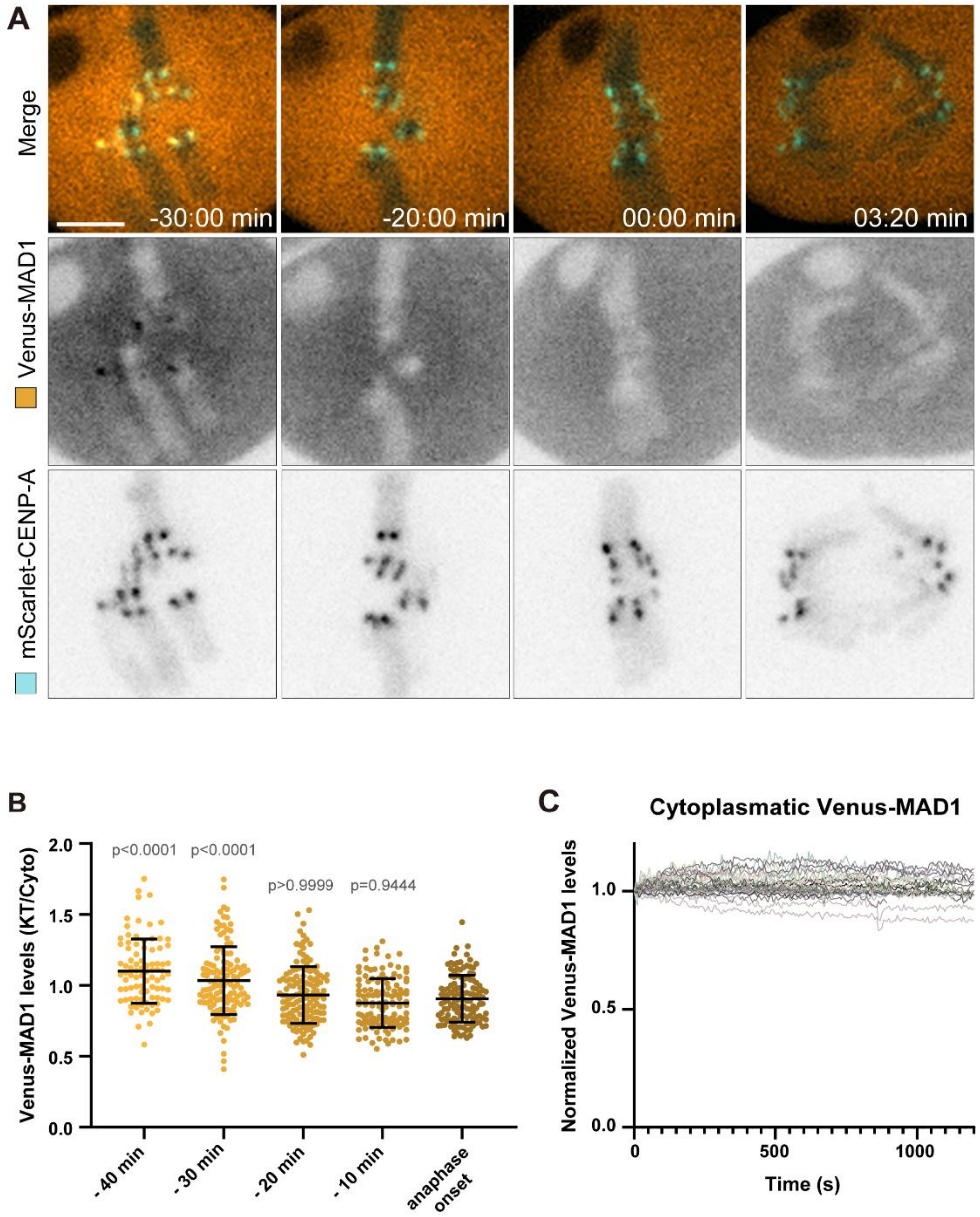


**Figure S1. MAD1 is undetectable at kinetochores 20 minutes before anaphase, related to Figure 1**

(A) Representative live-cell recording images of Indian muntjac fibroblasts expressing Venus-MAD1 (orange, upper panels or grey, lower panels) and mScarlet-CENP-A with SiR-tubulin labeled microtubules (grey, upper panels). Scale bar: 5 µm.

(B) Dot plot showing the ratio between kinetochore and cytoplasmatic Venus-MAD1 signal intensity at selected time points. Dots show individual kinetochores, lines show mean ±

s.d. (-40 min: 87 kinetochores, $p < 0.0001$; -30 min: 123 kinetochores, $p < 0.0001$; -20 min: 137 kinetochores, $p > 0.9999$; -10 min: 126 kinetochores, $p > 0.9444$; Anaphase onset: 126 kinetochores; 15 cells from 7 independent experiments)., P value calculated with Kruskal-Wallis test with Dunn’s comparisons of mean fluorescence in each time point with anaphase onset.

(C) Normalized signal intensity of cytoplasmatic Venus-MAD1 throughout live cell recording. Lines show selected cytoplasmatic regions (n= 27 spots, 9 cells, 6 independent experiments).

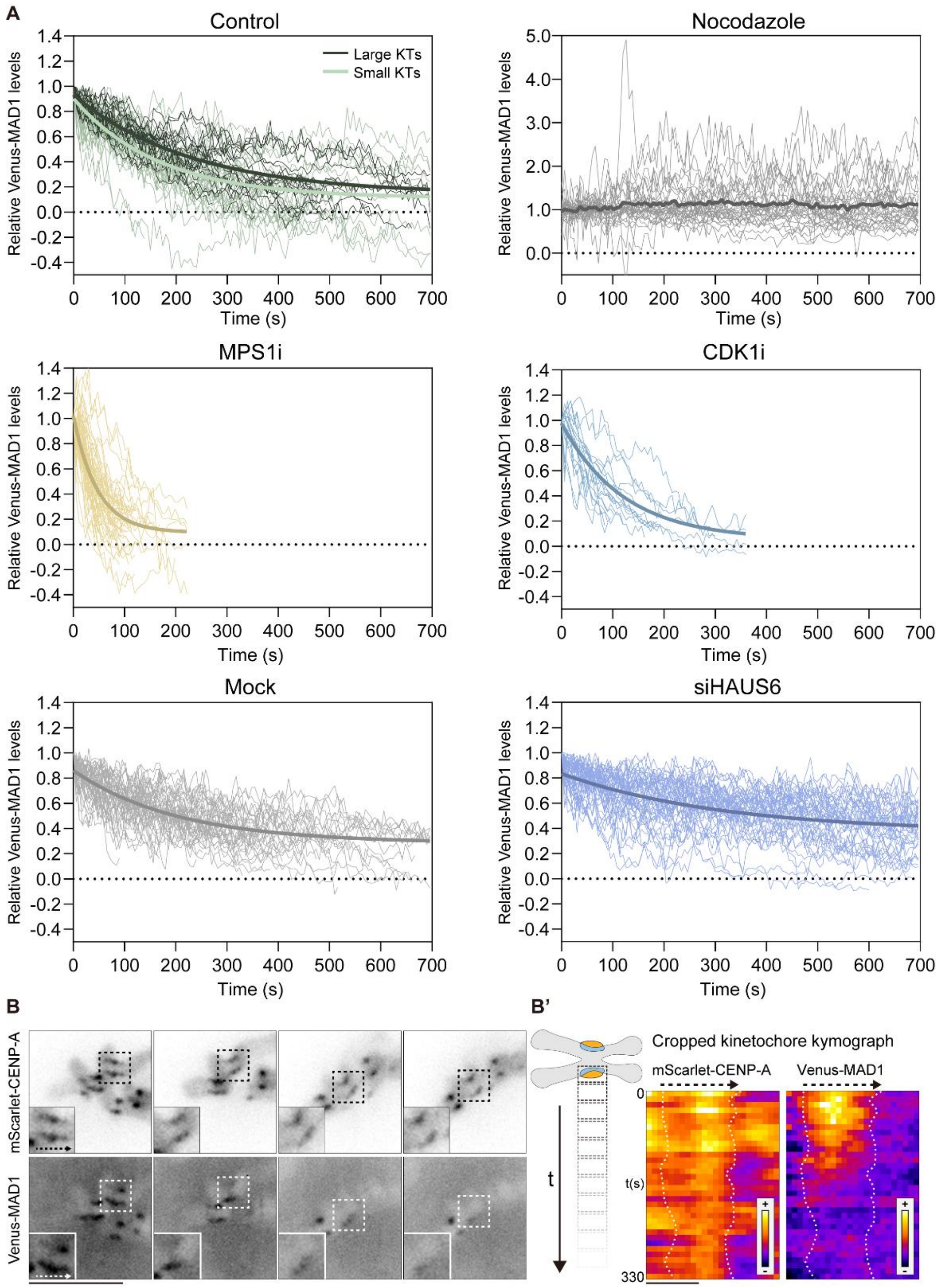


**Figure S2. MAD1 tracking on individual kinetochores throughout mitosis, related to Figure 1**

(A) Single kinetochore tracks of Venus-MAD1 fluorescence decay shown in Figure 1 and 6. Thin lines represent individual kinetochores (Control Large KTs n = 19 kinetochores, 7 cells, dark green; Control small KTs n = 34 kinetochores, 5 cells, light green; Nocodazole n = 31 kinetochores, 8 cells, grey; MPS1i n = 63 kinetochores, 12 cells, yellow; CDK1i n = 18 kinetochores, 10 cells, blue; Mock n = 48 kinetochores, 11 cells, light grey; siHAUS6 n = 47 kinetochores, 10 cells, purple blue; data pooled from at least four independent experiments per condition). Thick lines show the single-phase exponential decay fit curve for all pooled data (except for Nocodazole: average cure curve).

(B) Live-cell recording images of Indian muntjac fibroblasts stably co-expressing Venus-MAD1 (lower panel) and mScarlet-CENP-A (upper panel). Inset: kinetochore pair that shows non-uniform loss of MAD1 signal. Dashed arrow indicates the kinetochore depicted in B'. Scale bar: 5 μm.

(B') Kymographs depicting mScarlet-CENP-A (left) and Venus-MAD1(right) signal distribution along the longitudinal axis of the kinetochore shown in A throughout time. Signal intensity shown as a heatmap. Scale bar: 1 μm.

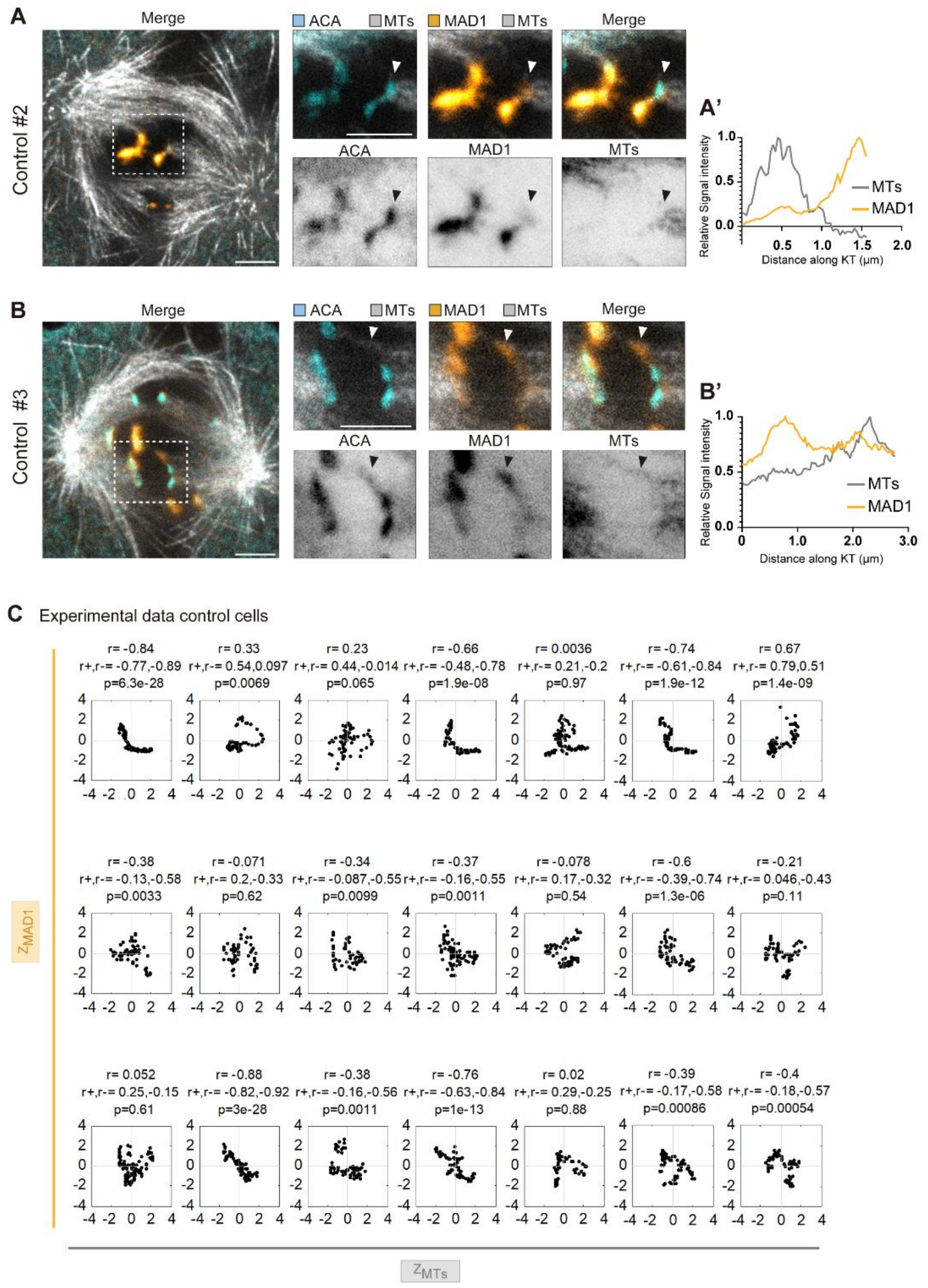


**Figure S3. Additional examples of partially attached kinetochores, related to Figure 3**

(A), (B) Representative CH-STED image of Indian muntjac fibroblasts with partially attached kinetochores. Microtubules (grey; STED), centromeres (ACA; blue; STED) and Venus-MAD1 (orange; confocal). Insets: partially attached kinetochores. Scale bar: 2 μm. (A'), (B') Max normalized Venus-Mad1 (orange) and tubulin (microtubules; grey) signal intensity profile along the longitudinal axis of the kinetochores highlighted in A and B, respectively (arrow head).

(C) Correlation map of MAD1 and microtubule intensity z-scores for the all partially attached kinetochores analyzed in untreated cells. Each scatterplot contains data from a single kinetochore, with each dot representing a sub-kinetochore region. P-value was calculated with Student t-test for zero correlation (two-sided).

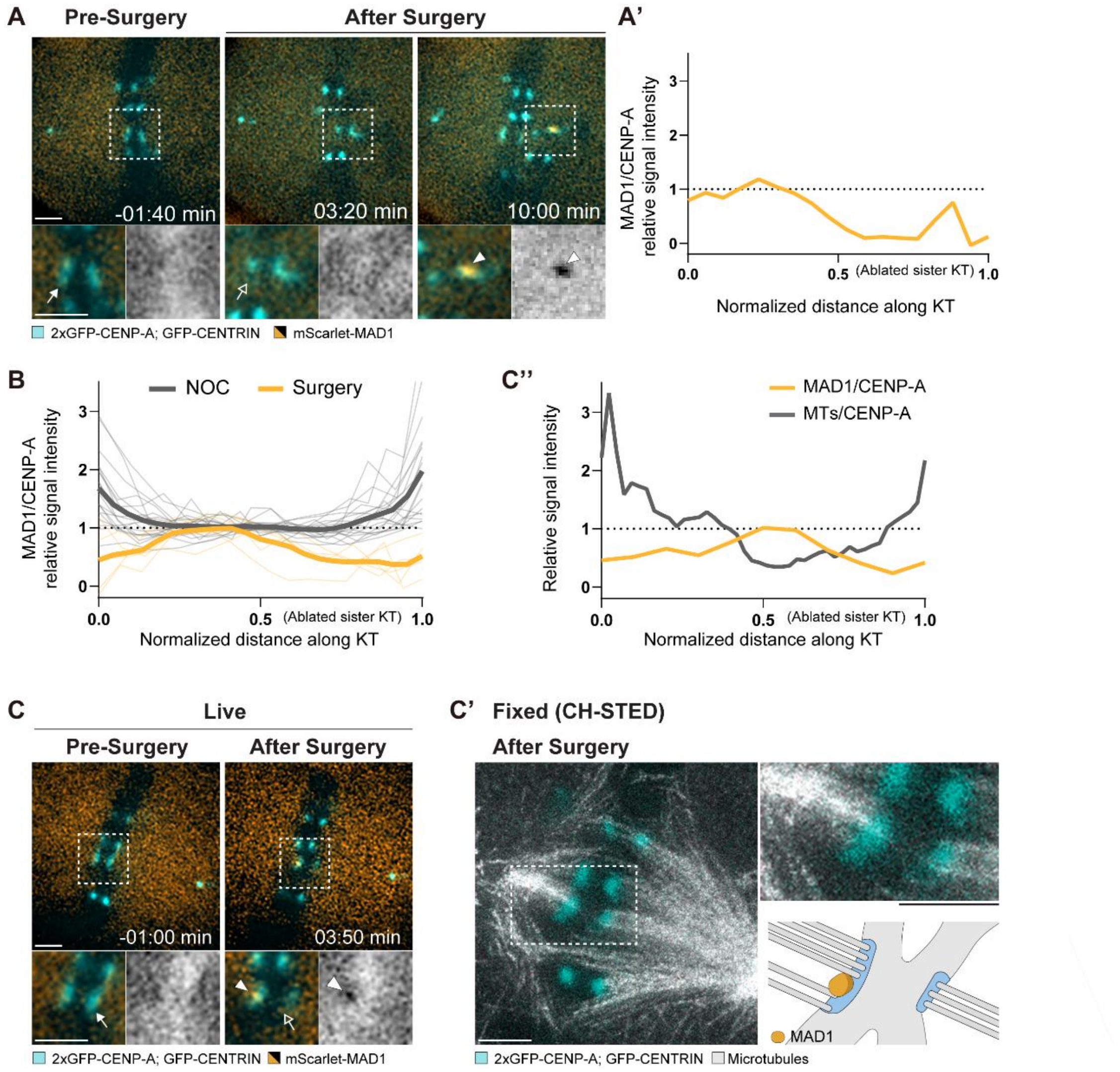


**Figure S4. Additional examples of localized MAD1 recruitment upon kinetochore laser ablation, related to Figure 4**

(A) Representative live-recording images of Indian muntjac cells expressing 2x-GFP-CENP-A and GFP-CENTRIN-1 (both in blue) and mScarlet-MAD1 (orange/ grey scale) upon partial kinetochore surgery. Insets: kinetochore pair that was perturbed. Arrows point to the ablated kinetochore region, pre- and post-surgery (solid and hollow arrow, respectively). Arrowhead shows MAD1 recruitment. Scale bar: 2 µm.

(A') Max normalized line intensity profile plot of MAD1/CENP-A signal ratio along the longitudinal axis of the kinetochore highlighted in A.

(B) Max normalized MAD1/CENP-A ratio intensity profile along the longitudinal axis of kinetochores after partial surgery of their sisters (orange) or upon acute Nocodazole treatment (NOC; grey). Thin lines show individual kinetochores and thick lines show average curves (surgery n = 5 kinetochores/cells, 5 independent experiments; Nocodazole n = 19 kinetochores/cells, 2 independent experiments). The graph depicts a cohort of cells that recruit MAD1 to a distal position relative to the surgery region (n = 5/16 MAD1 recruiting kinetochores). Data from Nocodazole treated cells is the same as shown in Figure 4.

(C), (C') Representative images from correlative live cell spinning disk (C) and fixed CH-STED microscopy (C') in a cell that recruited MAD1 to distal position relative to the surgery region. Indian muntjac cells expressing 2x-GFP-CENP-A and GFP-CENTRIN-1 (both in blue) and mScarlet-MAD1 (orange/grey scale) were fixed after partial kinetochore surgery and stained to visualize microtubules (grey). Insets: kinetochore pair that was perturbed. Arrows point to the ablated kinetochore region, pre- and post-surgery (solid and hollow arrow, respectively). Arrowheads show MAD1 recruitment. Scale bar: 2 μm.

(C'') Max normalized line intensity profile plot of MAD1/CENP-A (live cell data just before fixation, orange) and microtubules/CENP-A (fixed cell data, grey) signal ratio along the longitudinal axis of the kinetochore highlighted in C and C'.

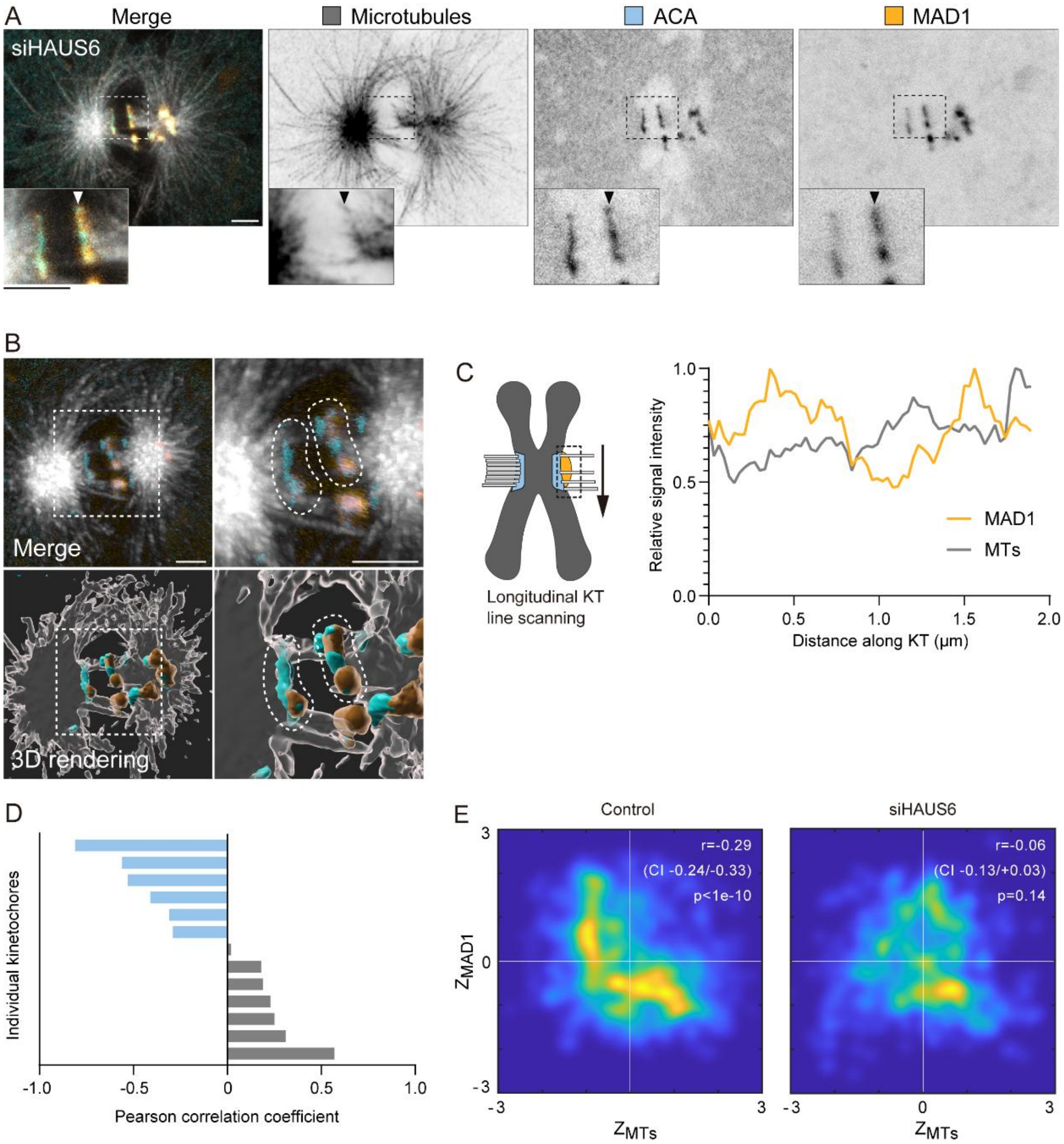


**Figure S5. Uniform perturbation of microtubule occupancy upon Augmin depletion, related to Figure 5**

(A) Representative CH-STED image of an Indian muntjac fibroblasts with partially attached kinetochores upon siHAUS6 treatment. Microtubules (grey; STED), centromeres (ACA; blue; STED) and Venus-MAD1 (orange; confocal). Insets: partially attached kinetochore. Scale bar: 2 μm.

(B) Three-dimensional reconstruction of the cell shown in A. Inset: partially attached kinetochore. The kinetochore pair of interest is traced by a dashed line. Microtubules (grey), centromeres (ACA; blue) and Venus-MAD1 (orange). Scale bars, Left: 3 μm; Right: 1 μm.

(C) Max normalized Venus-Mad1 (orange) and tubulin (microtubules; grey) signal intensity profile along the longitudinal axis of the kinetochore highlighted in A (arrow head).

(D) Pearson correlation coefficient between MAD1 and microtubules signal intensity along the kinetochore. Bars show individual partially attached kinetochores in siHAUS6 treated cells (n=13 kinetochores, 9 cells, 2 independent experiments).

(E) Correlation map of MAD1 and microtubule intensity z-scores for kinetochores in control (data from Figure 3) and in siHAUS6 treated cells (shown in D). The underlying scatterplot contains data from multiple kinetochores, with each dot representing a sub-kinetochore region. For visualization, the scatterplot was converted into a regional density map. P-value was calculated with Student t-test for zero correlation (two-sided).

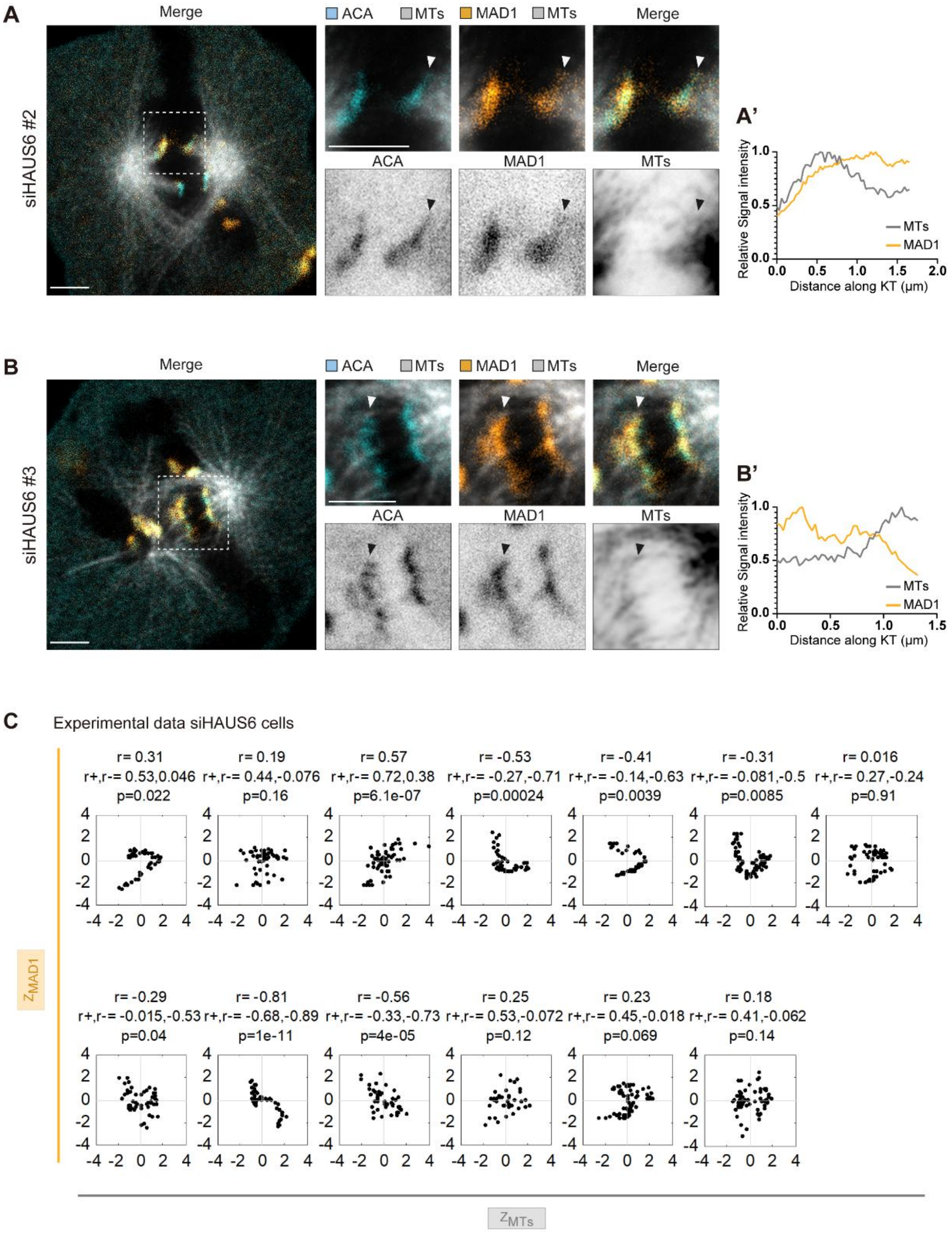


**Figure S6. Additional examples of partially attached kinetochores in Augmin depleted cells, related to Figure 5**

(A), (B) Representative CH-STED image of Indian muntjac fibroblasts siHAUS6 treated with partially attached kinetochores. Microtubules (grey; STED), centromeres (ACA; blue; STED) and Venus-MAD1 (orange; confocal). Insets: partially attached kinetochores. Scale bar: 2 µm.

(A'), (B') Max normalized Venus-Mad1 (orange) and tubulin (microtubules; grey) signal intensity profile along the longitudinal axis of the kinetochores highlighted in A and B, respectively (arrowhead).

(C) Correlation map of MAD1 and microtubule intensity z-scores for the all partially attached kinetochores analyzed in siHAUS6 cells. Each scatterplot contains data from a single kinetochore, with each dot representing a sub-kinetochore region. P-value was calculated with Student t-test for zero correlation (two-sided).

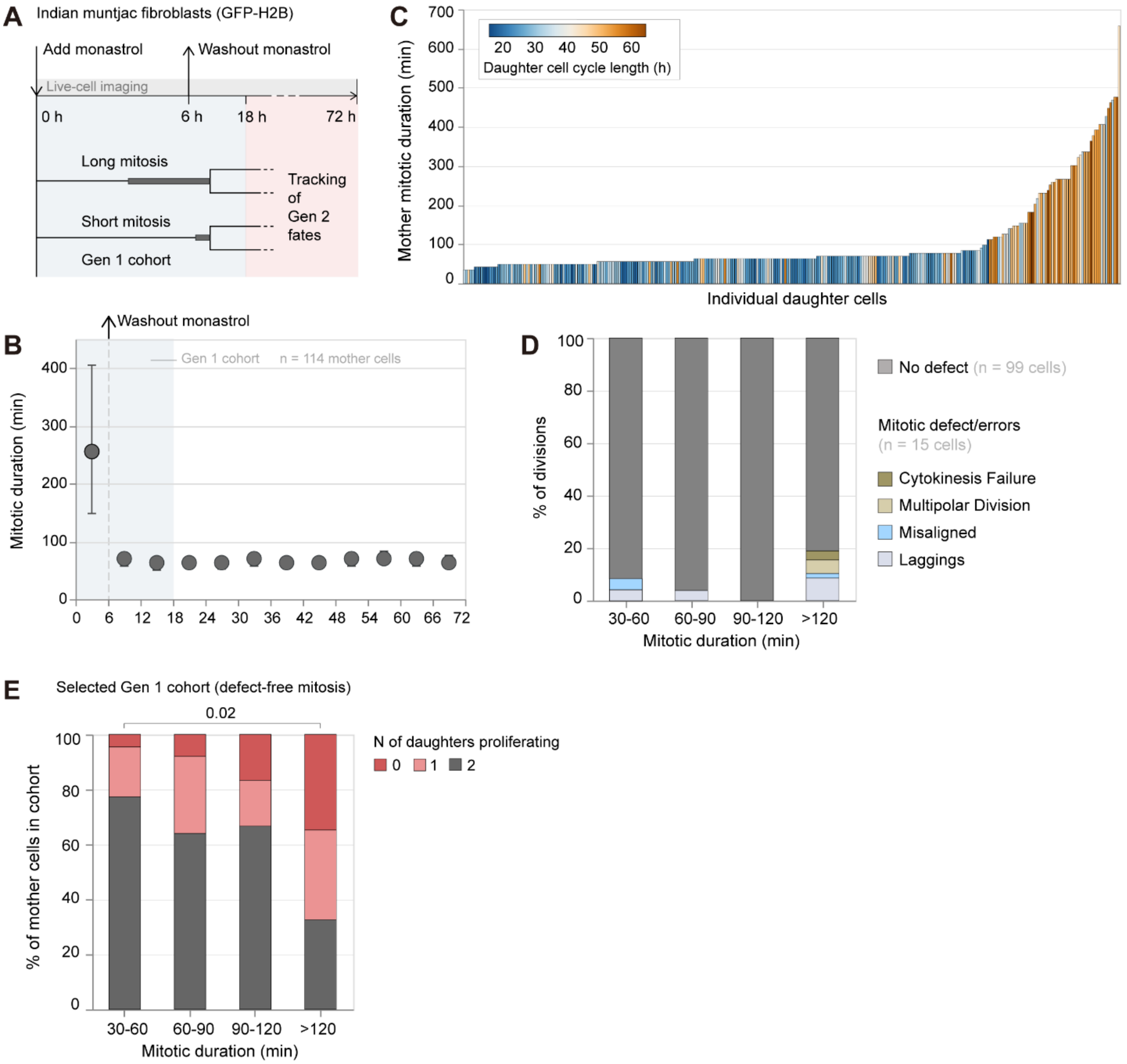


**Figure S7. Mitotic duration impacts daughter cell proliferation in Indian muntjac cells, related to Figure 6**

(A) Schematic summary of the live cell imaging setup and downstream analysis upon monastrol-induced arrest and release.

(B) Mitotic duration of Indian muntjac expressing H2B-GFP with increasing time after monastrol-induced arrest and release. Data pooled from two independent experiments (n = 492). Circles and error bars denote median and interquartile range in each time bin (bin size = 6 h).

(C) Daughter cell cycle duration as a function of mother mitotic duration from data shown in B. Bars show individual daughter cells (n = 305 cells pooled from two independent experiments). Note that only daughter cells that entered mitosis during imaging were included in the graph (see methods section).

(D) Bars show the percentage of mitosis that divide error-free or a with a particular mitotic defect/ error as a function of mitotic duration. Data shown in B filtered for cells that divide between 0 h to 18 h of recording. Data are binned (Gen 1 cohort; 30-60 min n = 24 cells, 60-90 min n = 26 cells, 90-120 min n = 6 cells, >120 min n = 58 cells, pooled from two independent experiments.

(E) Bars show the percentage of mother cells that has 0, 1 or 2 daughter cells dividing at least once, as a function of mother cells time in mitosis. Data filtered for cells that divide without defects within the selected Gen 1 cohort. Data are binned (30-60 min n = 22 cells, 60-90 min n = 25 cells, 90-120 min n = 6 cells, >120 min n = 46 cells, pooled from two independent experiments. 30-60 min vs >120 min, $p = 0.02$, chi-square test with Benjamini-Hochberg multiple comparison correction. For all other comparisons $p > 0.05$).
